# A foundation model learns the sequence and functional grammar of fully human heavy-chain-only antibodies

**DOI:** 10.64898/2026.09.10.750553

**Authors:** Nona Biosciences AI4S Team, Hongjiang Miao

**Affiliations:** Nona Biosciences (Shanghai) Co., Ltd., Shanghai, China.

## Abstract

Antibody language models learn from large natural repertoires, but whether generic representations capture the constraints of specialized antibody formats remains unclear. We first characterized fully human heavy-chain-only antibodies (HCAbs) independently of HCAb-trained models. Source-aware comparisons with conventional human VH domains revealed a reproducible distributional shift localized predominantly to CDR1/2, CDR3 architecture and, where supported, a restricted framework region rather than widespread framework remodeling. These model-independent differences motivated repertoire-specific pretraining. We developed HCAbLM, to our knowledge the first foundation model pretrained specifically on a large-scale fully human HCAb repertoire, using 31.8 million sequences from 73 independently immunized HCAb mice. HCAbLM learned a region-selective sequence-compatibility prior distinct from conventional antibody language models, and its frozen representations transferred to experimentally measured SEC purity, HIC behavior and thermal stability in grouped internal validation and retrospective cross-project evaluation. These findings identify repertoire composition as an important biological design variable for foundation models of specialized antibody formats.

## Introduction

Protein language models (PLMs) provide a general framework for learning representations of protein sequences without explicit structural or functional supervision. Trained on large collections of naturally occurring sequences, these models capture contextual dependencies among amino acids that reflect evolutionary and structural constraints and can transfer to tasks including structure prediction, mutational-effect prediction and functional annotation. The Evolutionary Scale Modeling (ESM) family established that sequence-only pretraining can recover biologically informative representations and that increasing model and data scale can improve their quality and generality [1]. These advances motivate the use of repertoire-scale pretraining to interrogate specialized protein sequence spaces.

Antibodies are particularly suited to this approach because immune-repertoire sequencing provides exceptionally large collections of naturally selected immunoglobulin sequences. The Observed Antibody Space (OAS) database enabled antibody-specific representation learning at scale [2], leading to models including AbLang [3], AntiBERTa [4], IgLM [5] and large paired antibody language models [6]. These models have supported sequence completion, contextual representation learning and conditional generation. More recent work has shown that antibody-specific objectives can also be used to address systematic properties of the training distribution, including germline bias [14]. Collectively, these studies demonstrate that the composition and design of antibody pretraining corpora can influence the representations learned by antibody language models.

Most antibody language models, however, have been trained predominantly on conventional immunoglobulin repertoires in which antigen recognition is mediated by paired heavy- and light-chain variable domains. It therefore remains unclear how well such representations capture sequence constraints imposed by alternative antibody architectures. Heavy-chain-only antibodies (HCAbs) provide a stringent test of this question because antigen recognition is mediated by a single variable domain in the absence of a conventional light-chain partner. HCAbs were first identified in camelids, where the VHH domain forms an autonomous antigen-binding unit [7]. Transgenic studies subsequently demonstrated that heavy-chain-only antibodies can be generated in mice and can undergo productive B-cell development and antigen-specific immune responses without conventional light-chain rearrangement [8].

Fully human HCAbs are especially informative in this context because they combine human immunoglobulin genetics with the structural requirements of a heavy-chain-only architecture. Transgenic mouse platforms carrying engineered human heavy-chain-only immunoglobulin loci have enabled the generation of soluble, antigen-specific and high-affinity fully human HCAbs [9,10]. In conventional antibodies, the heavy-chain variable domain (VH) and light-chain variable domain (VL) associate to form the variable fragment (Fv), whereas an HCAb variable domain must fold autonomously and form an antigen-binding surface without VL. Studies of camelid VHH domains and autonomous human VH domains have shown that residues at or near the former VH–VL interface can influence solubility, stability and autonomous folding [7,25–27]. Fully human HCAb repertoires may therefore occupy a specialized region of antibody sequence space shaped jointly by human germline constraints, heavy-chain-only structural requirements and in vivo selection.

These considerations raise a broader question: what aspects of a specialized antibody repertoire are captured by language-model pretraining? Sequence composition describes the prevalence of individual residues or motifs but does not capture higher-order dependencies among residues that determine whether an amino acid is compatible with its surrounding context. Masked or autoregressive language modeling estimates such conditional relationships directly [4,5]. We use the term sequence grammar to denote these contextual sequence constraints within a defined antibody repertoire. In this formulation, sequence grammar extends beyond marginal residue frequencies to include positional and combinatorial dependencies that characterize a molecular format.

Single-domain antibody-specific language modeling has precedent. VHHBERT was pretrained on approximately two million camelid VHH sequences, demonstrating that a repertoire restricted to a single-domain antibody format can support dedicated representation learning [11]. However, camelid VHHs and fully human HCAbs arise from different germline backgrounds and selection histories. Whether a large fully human HCAb repertoire encodes a distinct contextual sequence prior, and whether a foundation model trained specifically on that repertoire captures constraints that are incompletely represented by conventional antibody models, has not been systematically established.

A second question is whether repertoire-derived representations contain information relevant to molecular behavior beyond sequence reconstruction. Antibody developability reflects distributed sequence determinants that influence properties including hydrophobicity, aggregation, thermal stability, solubility and nonspecific interactions. Antibody language-model representations have been used to predict experimentally measured developability-related properties, including hydrophobic interaction chromatography and aggregation-associated measurements [12], and recent industrial-scale evaluations have assessed transfer to polyspecificity, HIC and self-interaction assays [13]. We therefore distinguish sequence grammar from functional grammar. Here, functional grammar is defined operationally as sequence-context information encoded by a pretrained representation that transfers to experimentally measured molecular properties. It does not imply that the model has learned antigen specificity, affinity or biological activity.

Here we ask whether the absence of a conventional light-chain partner is associated with a distinct repertoire-level sequence organization that motivates dedicated modeling. We first characterize functional fully human HCAbs and conventional human VH domains independently of any HCAb-trained representation, testing for whole-VH distributional differences under source-aware, near-duplicate-controlled evaluation and then localizing these differences through analyses of germline deviation, regional composition, CDR3 architecture and individual IMGT positions. These model-independent analyses establish the biological premise for repertoire-specific pretraining. We then train HCAbLM de novo on 31.8 million quality-controlled fully human HCAb sequences from 73 independently immunized HCAb mice and identify a performance–compute operating point for subsequent analyses. We next determine whether HCAbLM learns a region-selective sequence-compatibility prior distinct from those of conventional antibody language models and assess whether its frozen representations transfer to experimentally measured developability properties. Together, these analyses connect the molecular architecture of the heavy-chain-only format to repertoire-level sequence organization, repertoire-specific representation learning and experimentally relevant molecular-property prediction.

## Results

### Model-independent whole-VH analysis reveals a source-robust HCAb distributional shift

Because an HCAb variable domain must fold and engage antigen without a conventional VL partner, both the former VH–VL interface and the antigen-binding loops operate in a distinct structural context (Fig. 1a,b). We therefore first asked, independent of any HCAb-trained representation, whether functional fully human HCAbs occupy a reproducibly different sequence distribution from conventional human VH domains.

**Fig 1.**
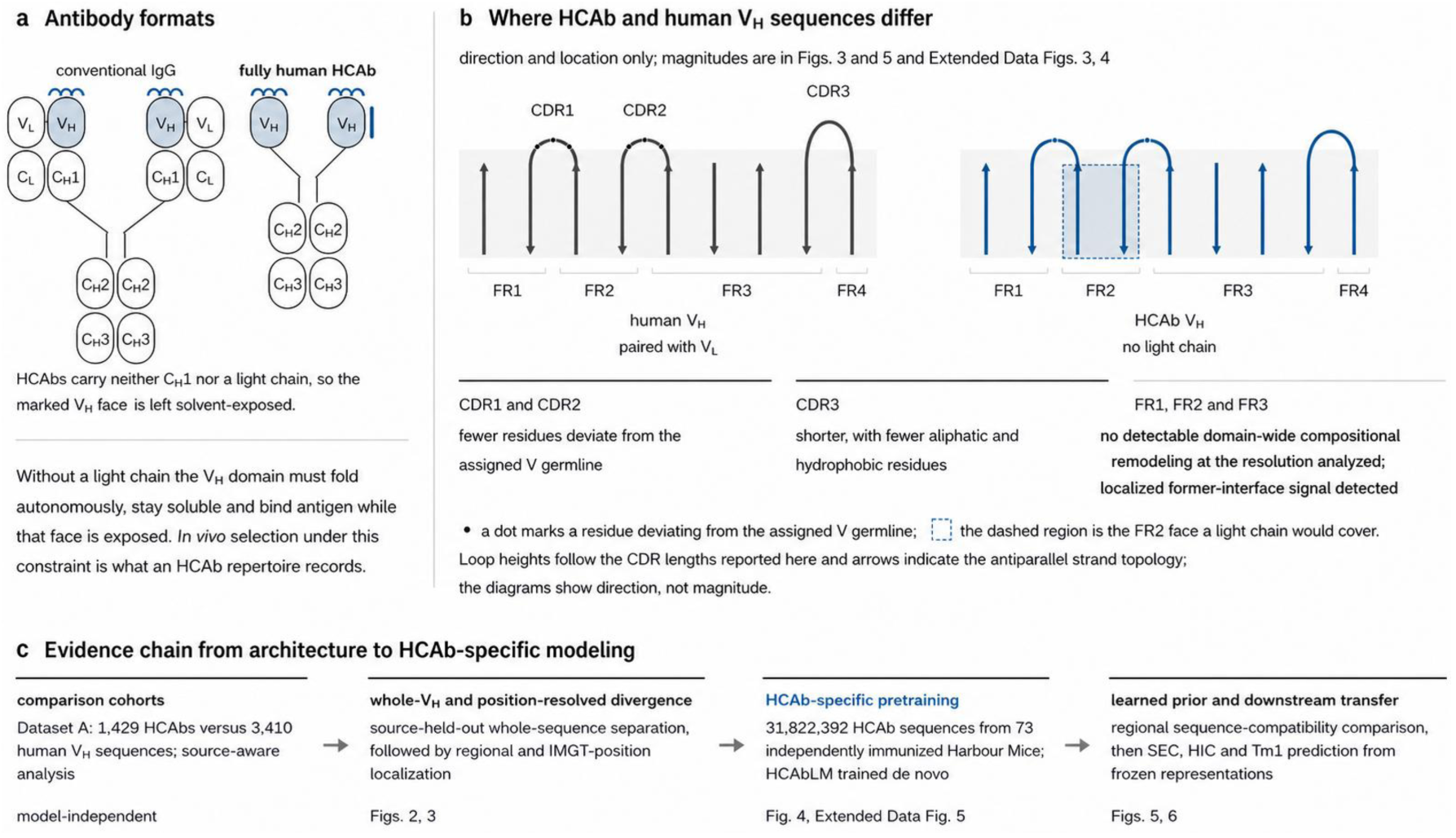
Molecular premise and evidence chain. **a,** A conventional IgG pairs a heavy-chain variable domain (VH) with a light-chain variable domain (VL), whereas a fully human HCAb lacks CH1 and a light chain, leaving the surface normally contacted by VL solvent-exposed. **b,** Schematic topology of the compared variable domains in IMGT framework (FR1–FR4) and complementarity-determining (CDR1– CDR3) regions. Dots indicate positions differing from the assigned V germline; the dashed region marks the light-chain-facing framework surface. Schematics indicate direction and location rather than effect magnitude and are not structural measurements. **c,** Study workflow progressing from source-aware, model-independent HCAb–human VH comparison through whole-VH discrimination and position-resolved localization, followed by HCAb-specific pretraining, regional learned-prior comparison and developability evaluation. HCAbs showed lower CDR1/2 NGL burden and shorter, less hydrophobic CDR3s; no detectable domain-wide framework remodeling was observed, whereas targeted analysis identified a localized former VH–VL-interface signal. Quantitative effects and uncertainty are reported in Figs. 2–6, Extended Data Figs. 1–5 and Source Data.

We performed a cross-fitted whole-VH kernel analysis on Dataset A, restricting the primary comparison to five shared IGHV genes represented by at least five independent sources in each cohort (IGHV3-11, IGHV3-23, IGHV3-33, IGHV3-53 and IGHV3-74). After removing 35 sequences belonging to cross-cohort near-duplicate components, the analysis included 1,400 HCAbs from 17 primary sources and 1,545 human VH sequences from 184 primary sources. The prespecified kernel integrated complementary whole-sequence information from pooled IMGT-aligned residue and gap states and complete-sequence 2-mer and 3-mer spectra, without using HCAbLM or any other HCAb-trained embedding.

In fivefold source-held-out evaluation, all sequences from a given source were assigned to the same fold, and training sequences belonging to a near-duplicate component shared with a test sequence were removed within each fold. The equal- IGHV raw-kernel cross-fitted witness separation was 0.0204 (95% source-bootstrap CI, 0.0177–0.0231; family-global Bonferroni P ≤ 0.001), and each of the five IGHV-specific permutation tests remained significant after Holm adjustment (Fig. 2a). On the standardized out-of-fold scale used for visualization, the source- and IGHV-equal weighted mean witness score was -1.40 for human VH and +0.88 for HCAb. The separation was consistent across sources: all 17 HCAb sources had positive mean witness scores, whereas 176 of 184 human VH sources had negative mean scores (Fig. 2b). Thus, the whole-VH signal was not driven by a small subset of disproportionately large sources.

**Fig 2.**
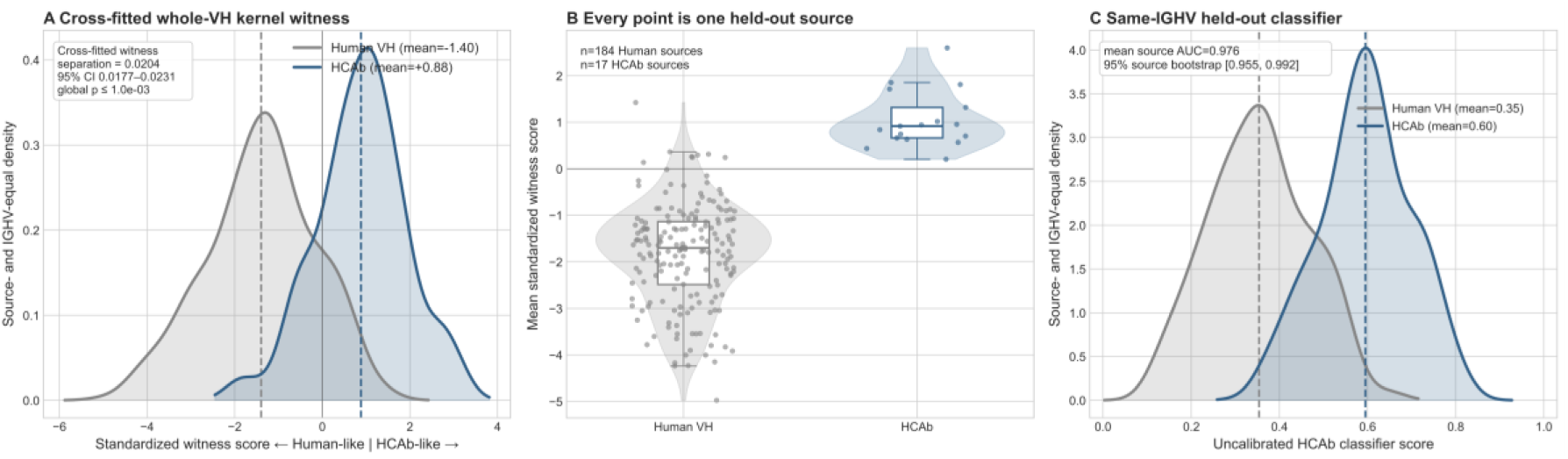
Functional HCAbs and human VH domains occupy distinct whole-sequence distributions within shared IGHV strata. The analysis included 1,400 HCAbs from 17 primary sources and 1,545 human VH sequences from 184 primary sources across five shared IGHV genes (IGHV3-11, IGHV3-23, IGHV3-33, IGHV3-53 and IGHV3-74). Entire sources were held out in fivefold cross-fitting, and training sequences sharing a frozen near-duplicate component with a test sequence were removed within each fold. **a,** Standardized out-of-fold witness scores from a prespecified whole-VH kernel combining IMGT-aligned residue/gap states with complete-sequence 2-mer and 3-mer cosine similarities. Dashed lines indicate source- and IGHV-equal cohort means (human VH, -1.40; HCAb, +0.88). The equal-IGHV cross-fitted witness separation is 0.0204 in raw kernel units (5,000-replicate source-bootstrap 95% CI, 0.0177-0.0231; Bonferroni global-null bound, *P* ≤ 0.001); all five IGHV-specific permutation tests remained significant after Holm adjustment. **b,** Mean standardized out-of-fold witness score for each held-out primary source. Points represent sources; violins show source-level distributions and boxes indicate medians and interquartile ranges. All 17 HCAb sources had positive mean scores, whereas 176 of 184 human VH sources had negative mean scores. **c,** Out-of-fold scores from IGHV-specific L2-regularized logistic classifiers evaluated using the same source-held-out splits and near-duplicate exclusion procedure. The equal-IGHV mean source-level AUROC was 0.976 (95% source-bootstrap CI, 0.955– 0.992); dashed lines indicate mean classifier scores (human VH, 0.35; HCAb, 0.60). Classifier outputs were uncalibrated ranking scores and were used for discrimination rather than probability estimation. Density curves in **a** and **c** give equal total weight to each IGHV gene, each source within IGHV and each sequence within source. No HCAb-trained representation was used in these analyses.

We next asked whether the same distinction could be recovered using a complementary discriminative formulation. IGHV-specific L2-regularized logistic classifiers were trained using the same whole-sequence inputs, source-held-out folds and near-duplicate exclusion scheme. The equal-IGHV mean source-level AUROC reached 0.976 (95% source-bootstrap CI, 0.955–0.992; Fig. 2c), with mean uncalibrated classifier scores of 0.35 for human VH and 0.60 for HCAb. Together, the kernel and classifier analyses demonstrate a reproducible whole-VH distributional distinction between functional HCAbs and conventional human VH domains within Dataset A, independent of HCAb-specific model representations. These analyses establish the existence and source-level reproducibility of the sequence shift, while its regional organization and structural interpretation are examined below.

### Position-resolved repertoire analysis localizes HCAb–human VH sequence differences

The whole-VH analysis established a reproducible source-level distributional distinction between functional HCAbs and conventional human VH domains but did not resolve the sequence features underlying that separation. We therefore localized the shift using complementary analyses of CDR3 architecture, regional germline deviation, amino-acid composition and IMGT-position-resolved sequence variation.

For these analyses, we compared 1,429 mature functional HCAbs with 3,410 publicly available assay-positive human VH sequences in Dataset A, with equal total weight assigned to 17 HCAb and 360 human source units. Human comparators were restricted to the same IGHV gene set represented by the transgenic platform. HCAb CDR3s were shorter than human VH CDR3s (mean, 13.20 versus 15.16 amino acids; difference, -1.96 amino acids; 95% CI, -2.72 to -1.20; q < 0.0001) and remained shorter after equal weighting across shared IGHV strata (-1.77 amino acids). Across reliably mapped V-derived positions, the non-germline-like (NGL) fraction was lower in HCAbs in CDR1 (8.05% versus 16.52%; difference, -8.47 percentage points; q < 0.0001) and CDR2 (12.32% versus 17.34%; difference, -5.02 percentage points; q = 0.0013), whereas no difference was detected for the pooled framework (3.48% versus 5.05%; difference, -1.57 percentage points; q = 0.17) (Extended Data Fig. 3). Thus, the pooled framework analysis did not support broad compositional remodeling, although it did not establish framework equivalence.

Regional amino-acid composition showed a concordant, predominantly CDR- centered pattern. HCAb CDR1 was enriched in aromatic residues and depleted in aliphatic/hydrophobic residues, CDR2 was enriched in polar uncharged residues, and CDR3 was depleted in aliphatic/hydrophobic residues (Extended Data Fig. 3). Amino-acid composition and substitution effects meeting the prespecified false-discovery criterion were confined to CDR1/2 (Extended Data Fig. 4). Equal weighting across shared IGHV strata attenuated several marginal CDR1/2 composition effects and eliminated the proportional CDR1-versus-framework contrast, indicating that IGHV usage contributes to part of the regional signal. By contrast, shorter CDR3s, reduced CDR3 hydrophobicity and lower CDR1/2 NGL fractions were retained after reweighting. These analyses therefore support a regionally organized HCAb sequence distribution dominated by changes in the antigen-binding loops rather than uniform domain-wide remodeling.

The pooled framework, however, masked a localized exception at the former VH–VL interface. In a prespecified partition of V-derived framework positions, four hallmark positions at this interface (IMGT 42, 49, 50 and 52) showed a higher NGL fraction in HCAbs than in human VH sequences (5.78% versus 1.38%; difference, +4.40 percentage points; 95% CI, +1.70 to +7.83; q < 0.001), whereas the remaining framework showed the opposite direction (3.36% versus 5.24%; difference, -1.88 percentage points; 95% CI, -2.70 to -1.13; q < 0.001). The resulting patch-by-cohort interaction was +6.28 percentage points (95% CI, +3.77 to +9.51; q < 0.001), remained essentially unchanged after equal weighting across shared IGHV genes (+6.27 percentage points), and showed no direction reversal in leave-one-HCAb- source-out analysis. In an exploratory 96-position profile, the HCAb-enriched signal within this patch localized most strongly to IMGT 52 (+6.39 percentage points; pointwise 95% CI, +1.60 to +13.74; q = 0.006), whereas IMGT 42, 49 and 50 did not individually meet the detection criterion and the simultaneous interval for IMGT 52 included zero (Fig. 3). At the residue-class level, the hallmark patch was also less hydrophobically enriched relative to the remaining framework in HCAbs than in human VH sequences (interaction, -2.32 percentage points; pointwise 95% CI, -4.22 to -0.96; simultaneous 95% CI, -4.53 to -0.12; q = 0.001), with the same direction retained after equal weighting across shared IGHV genes (-3.66 percentage points). Thus, HCAbs do not show uniform framework remodeling, but the absence of a pooled framework difference conceals a localized sequence shift at the former VH– VL interface.

**Fig. 3.**
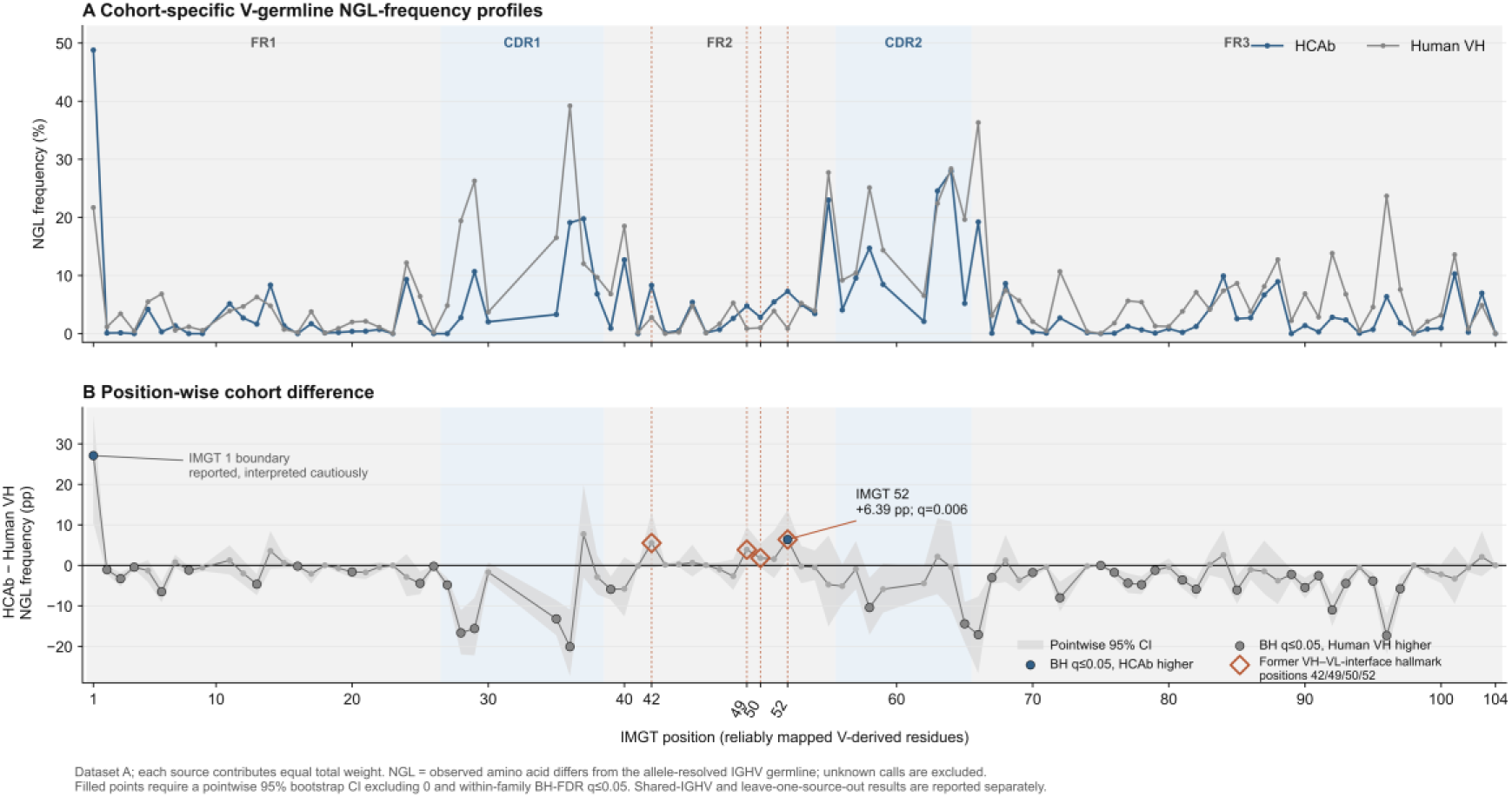
Position-resolved V-germline-deviation profiles reveal a localized signal at the former VH–VL interface. **a,** Source-equal NGL-frequency profiles for Dataset A, comprising 1,429 HCAbs and 3,410 human VH sequences from 17 and 360 source units, respectively, across reliably mapped V-derived integer IMGT positions. **b,** Source-equal HCAb–human VH differences in NGL frequency across 96 callable positions; Shaded band indicates pointwise 95% confidence intervals from two-stage bootstrap resampling. Filled points denote positions for which the pointwise interval excluded zero and the within-family Benjamini–Hochberg-adjusted q value was ≤0.05. Orange vertical lines and diamonds mark the prespecified former VH– VL-interface hallmark positions IMGT 42, 49, 50 and 52; these annotations do not indicate that all four positions were individually significant. Within this patch, the HCAb-enriched signal was strongest at IMGT 52 (+6.39 percentage points; q = 0.006), although its simultaneous family-wise interval included zero, and it is therefore interpreted as a localization signal rather than a confirmatory single-position effect. The signal at IMGT position 1 is shown for completeness but interpreted cautiously because of its proximity to the N-terminal annotation boundary. Shared-IGHV weighting and leave-one-HCAb-source-out sensitivity analyses are reported separately in the Source Data.

Together, the whole-sequence and position-resolved analyses demonstrate that functional HCAbs and conventional human VH domains occupy reproducibly distinct sequence distributions within Dataset A. The differences were regionally organized, with the strongest effects in CDR1/2 and CDR3 together with a localized shift at the former VH–VL interface, rather than widespread framework remodeling. These model-independent observations provide the biological rationale for asking whether a foundation model pretrained directly on the fully human HCAb repertoire can learn this region-selective sequence organization.

### HCAb-specific pretraining and model scaling identify HCAbLM-300M as an efficient operating point

Motivated by these model-independent sequence differences, we assembled a unified HCAb pretraining corpus comprising 31,822,392 model-eligible, globally unique HCAb VH amino-acid sequences from 73 independently immunized Harbour Mice spanning eight distinct antigens. Descriptive repertoire annotation showed concentrated IGHV and IGHJ usage and an annotation-derived CDR3-length distribution centered at 12–14 amino acids (Extended Data Fig. 5). V(D)J assignments and V/J germline alignments were used to classify reliably mapped residues as germline matched or non-germline-like (NGL), whereas true CDR3/junctional and incompletely resolved positions were treated separately. During self-supervised pretraining, the fixed 15% masking budget preferentially sampled true CDR3 and high-confidence NGL positions, and model optimization used a prespecified region- balanced focal-loss objective to emphasize these biologically informative sequence classes (Fig. 4a; Methods).

**Fig. 4.**
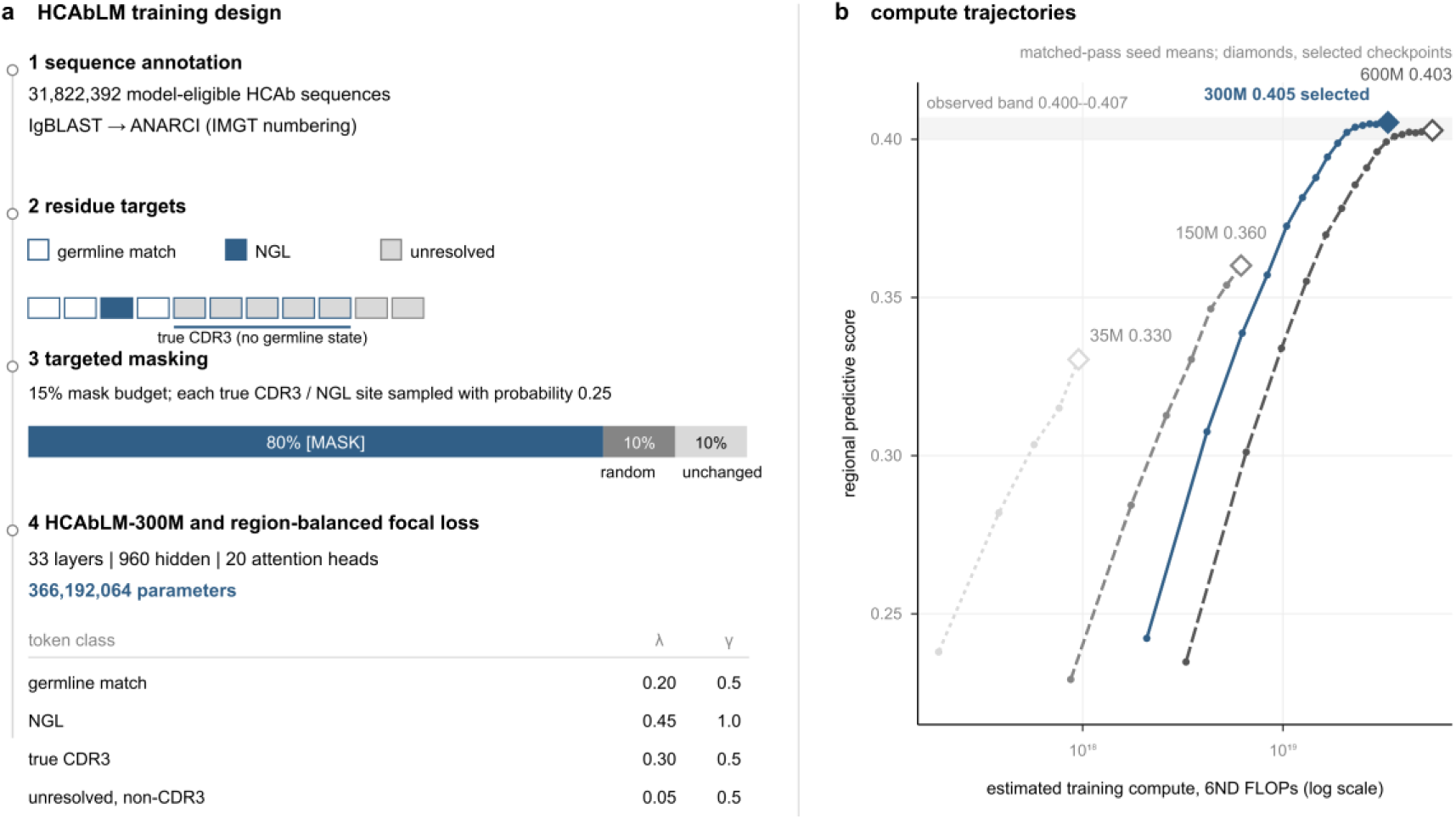
HCAbLM training design and performance–compute operating-point selection. **a,** V(D)J annotation and V/J-germline alignments were used to classify reliably mapped residues as germline matched, non-germline-like (NGL) or unresolved, with true CDR3 positions defined independently from sequence coordinates. A fixed 15% masking budget preferentially sampled true CDR3 and high-confidence NGL positions, followed by 80:10:10 mask/random/unchanged corruption. HCAbLM-300M is a de novo ESMC-style encoder-only Transformer with 366.19 million parameters trained using a region-balanced focal-loss objective. **b,** Measured checkpoint trajectories as a function of estimated training compute. The regional predictive score was defined as exp[-(0.60 × PCE_NGL_ + 0.40 × PCE_CDR3_)], where higher values indicate lower region-weighted prediction loss. Performance was evaluated on a frozen 80,000-target diagnostic bank with inverse-probability weighting to recover the natural positional mixture. Lines connect measured checkpoints and are not fitted curves; where matched-pass replicate seeds were available, plotted coordinates show their mean. Diamonds indicate checkpoints selected using full-sequence validation, and the shaded band denotes the observed score range of 0.400–0.407. HCAbLM-300M reached this range with lower estimated compute than the 600M configuration and was selected as the performance– compute operating point among evaluated configurations. Model-size and checkpoint selection used only the prespecified validation panel; Dataset A, Dataset B and downstream developability outcomes were not used for selection.

We trained ESMC-style encoder-only Transformers de novo in four nominal size classes spanning 35M to 600M parameters [15]. Model scaling was evaluated on a frozen 80,000-target diagnostic bank using a region-weighted predictive score that integrates performance at high-confidence NGL and true-CDR3 positions, with higher values indicating better prediction. Across training trajectories, the score increased with compute and approached an observed plateau of approximately 0.400–0.407 (Fig. 4b). HCAbLM-300M, containing 366.19 million parameters, reached this performance band with less estimated training compute than the 600M configuration and achieved the lowest mean region-weighted loss in full-sequence validation. We therefore selected HCAbLM-300M as the performance–compute operating point for all subsequent analyses. Where matched-pass replicate seeds were available, Fig. 4b shows their mean trajectories; diamonds denote the checkpoints selected by the frozen validation procedure. Model-size and checkpoint selection relied exclusively on the prespecified validation panel; Dataset A, Dataset B and developability outcomes were not used for model selection.

### HCAbLM captures a region-selective compatibility prior distinct from conventional antibody models

Having selected HCAbLM-300M as the performance–compute operating point, we next asked whether repertoire-specific pretraining altered the contextual sequence preferences learned across the HCAb variable domains. We measured per-residue prediction loss in Dataset A using the frozen HCAbLM-300M checkpoint together with IgLM [5] and AbLang-2 [14]. Because these models use different scoring formulations, raw losses were retained on their native scales and cross-model effects were compared only after within-model standardization by source-level variation (Methods). For reference, the model-independent analysis showed substantially lower NGL fractions in HCAb CDR1/2 but no detectable difference when the framework was pooled (Fig. 5a), providing a compositional context for the model- based comparisons.

**Fig. 5.**
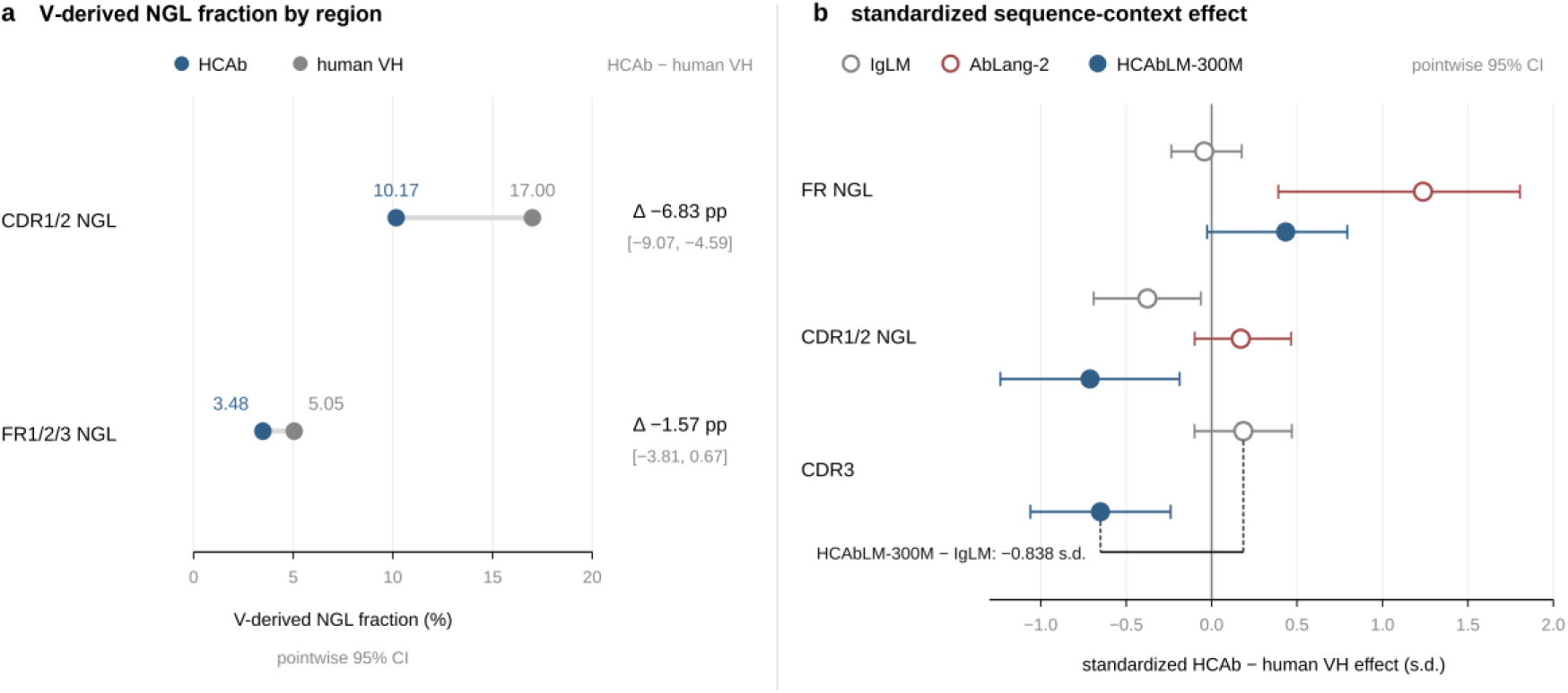
HCAbLM reveals a region-selective sequence-compatibility prior relative to model-independent repertoire differences. Dataset A comprises 1,429 mature functional HCAbs and 3,410 publicly available assay-positive human VH sequences, representing 17 and 360 source units, respectively, with both cohorts restricted to the shared IGHV gene set. **a,** Source-equal NGL fractions at reliably mapped V-derived positions in CDR1/2 and the pooled framework. Brackets indicate 95% bootstrap confidence intervals for HCAb–human VH differences. **b,** Within-model standardized HCAb–human VH effects on per-residue prediction loss for framework NGL residues, CDR1/2 NGL residues and CDR3 under IgLM, AbLang-2 and HCAbLM-300M. Negative values indicate lower prediction loss for HCAbs under the indicated model. Raw prediction scores were retained on their native model-specific scales and were not compared directly across models; displayed effect sizes were standardized using source-level variation. CDR3 was analyzed separately because germline status was defined only for reliably mapped V-derived positions. Pointwise 95% confidence intervals and Benjamini–Hochberg-adjusted q values were evaluated within prespecified analysis families; corresponding family-wise simultaneous intervals are reported in the Source Data. Full regional decompositions and model-independent composition analyses are shown in Extended Data Figs. 1–4. HCAbLM-300M denotes the frozen checkpoint selected as described in Methods.

The clearest model-dependent divergence occurred in CDR3. HCAbLM-300M assigned lower standardized prediction loss to HCAb than to human VH CDR3 residues (HCAb - human VH, -0.651 s.d.; 95% CI, -0.959 to -0.343), whereas IgLM assigned higher loss to HCAb CDR3s (+0.186 s.d.), yielding a paired HCAbLM–IgLM contrast of -0.838 s.d. (95% CI, -1.075 to -0.649) (Fig. 5b). AbLang-2 showed the same direction as IgLM. Equal weighting across the 18 CDR3-length strata shared by the two cohorts preserved these opposing directions, indicating that the difference was not explained by the shorter CDR3 distribution of HCAbs alone.

Regional effects outside CDR3 were more heterogeneous. Both external antibody models assigned lower prediction loss to HCAb CDR1 residues, whereas a CDR2 difference was detected only by HCAbLM-300M. Among germline-deviating framework residues, AbLang-2 assigned markedly higher loss to HCAbs (+1.237 s.d.; 95% CI, +0.391 to +1.805; q = 0.0020), despite the absence of a detectable difference in the pooled framework NGL fraction; HCAbLM-300M showed the same direction without meeting the prespecified detection criterion, whereas IgLM showed little difference. FR3 also showed a diffuse region-level effect under the external models, but the corresponding simultaneous family-wise interval included zero and no individual position met the confirmatory threshold. These observations further distinguish compositional frequency from contextual compatibility: regions with similar aggregate germline-deviation burdens can nevertheless differ in the sequence contexts preferred by a language model.

Dataset B, comprising 232 HCAbs and 232 IGHV-matched conventional human VH sequences, provided an orthogonal provenance sensitivity analysis. Regional prediction-loss effects were again non-uniform across the variable domain, although the direction of individual regional and whole-domain contrasts was not identical to Dataset A (Extended Data Figs. 1 and 2). Together, these analyses show that HCAbLM does not simply assign a uniformly higher compatibility to HCAb sequences. Instead, repertoire-specific pretraining is associated with a redistribution of contextual compatibility across sequence regions, most prominently in CDR3. The opposing CDR3 preferences of HCAbLM and conventional antibody language models are consistent with an important contribution from HCAb-specific repertoire pretraining, while differences in model architecture, training objective, scoring formulation and prior sequence exposure prevent attribution to training-corpus composition alone.

### HCAbLM representations retain developability-associated information across grouped validation and retrospective cross-project benchmarks

To determine whether the learned sequence representations captured information relevant to experimentally measured molecular properties, we froze each encoder and trained shallow endpoint-specific prediction heads for size-exclusion chromatography (SEC) purity, hydrophobic interaction chromatography (HIC) behavior and the first thermal transition midpoint (Tm1). All benchmark sequences and their identified clonal relatives were excluded from HCAbLM pretraining, and related sequences were assigned to the same cross-validation group to limit performance inflation from sequence similarity (Methods).

In grouped nested cross-validation, different representations performed best for different developability endpoints. HCAbLM-300M achieved an AUROC of 0.691 for the internally defined SEC classification endpoint (1,469 sequences in 773 related- sequence groups). HCAb-adapted IgBERT [6] yielded the highest HIC correlation (Spearman’s ρ = 0.536; 506 sequences in 246 groups), whereas ESMC-6B [15] yielded the highest Tm1 correlation (ρ = 0.430; 425 sequences in 209 groups). Thus, HCAb- specific pretraining did not confer uniform superiority across properties, but its frozen representation retained predictive information for a chromatographic developability endpoint without task-specific encoder fine-tuning.

We next assessed transfer across antibody projects using the same frozen encoders and prediction heads without refitting on the project-level evaluation data (Fig. 6). HCAbLM-300M gave the highest project-macro point estimate for SEC purity (ρ = 0.355; 87 sequences from two projects) and HIC favorability (ρ = 0.453; 108 sequences from three projects), whereas ESMC-6B gave the highest point estimate for Tm1 (ρ = 0.272; 108 sequences from three projects). For SEC, the internal benchmark used the prespecified binary purity endpoint, whereas cross-project transfer was evaluated against continuous SEC purity using Spearman correlation (Methods). Because performance on these held-out projects contributed to endpoint-level model comparison, the cross-project estimates are selection-affected and are interpreted as retrospective benchmarking rather than independent validation. Collectively, these analyses show that frozen HCAbLM representations contain developability-associated information that transfers across sequence groups and antibody projects, while the relative advantage over alternative protein and antibody encoders remains endpoint dependent and requires confirmation in a prospectively locked evaluation set.

**Fig. 6.**
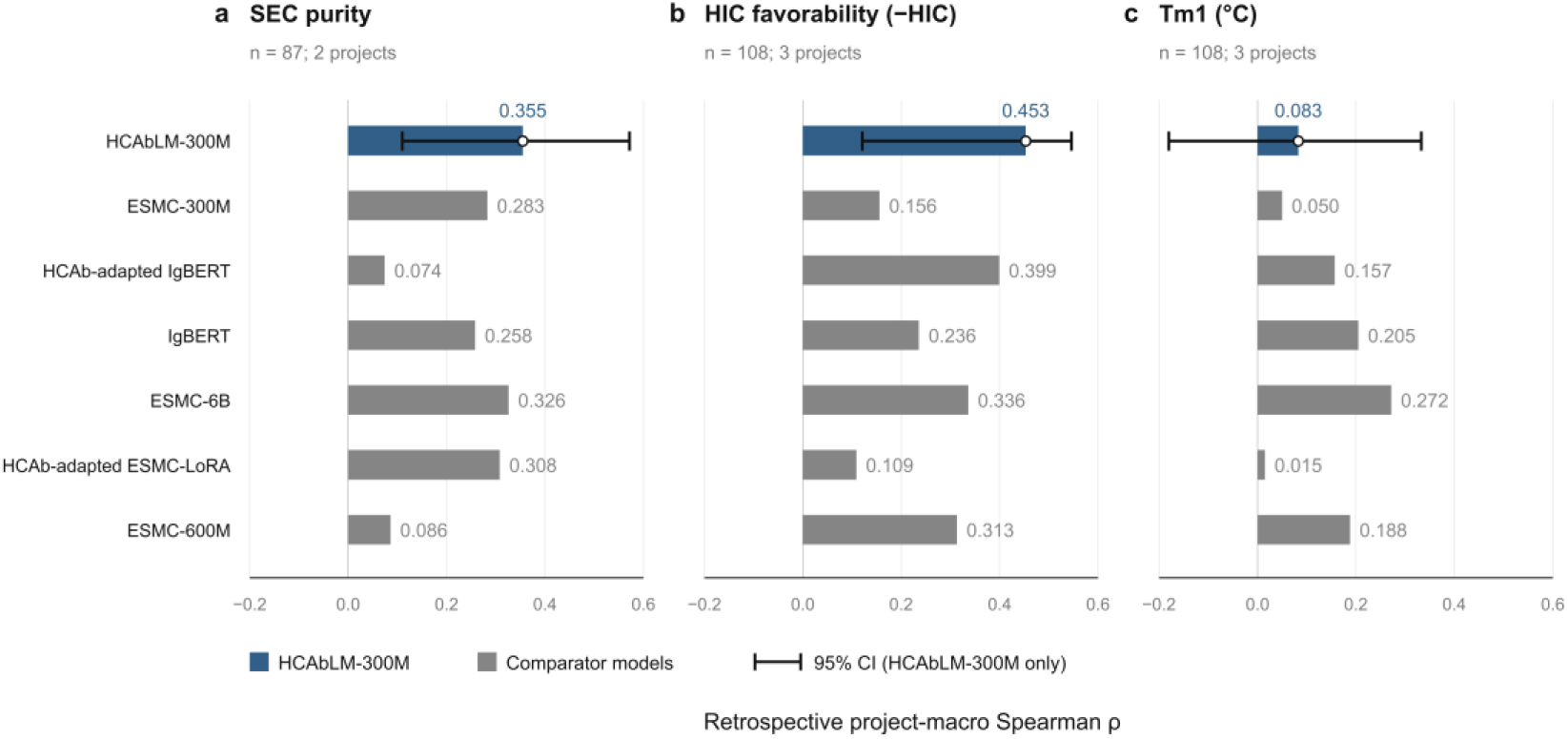
Frozen sequence representations transfer to held-out developability projects. Seven candidate encoders are compared for **a,** SEC purity, **b,** HIC favorability and **c,** Tm1 using retrospective project-macro Spearman’s ρ. SEC includes 87 sequences from two projects; HIC and Tm1 each include 108 sequences from three projects. Prediction heads were fitted on internal data and were not refitted on the held-out projects. HCAbLM-300M gave the highest point estimates for SEC (ρ = 0.355) and HIC (ρ = 0.453), whereas ESMC-6B gave the highest point estimate for Tm1 (ρ = 0.272).

## Discussion

This study links the molecular architecture of fully human heavy-chain-only antibodies to repertoire organization, repertoire-specific representation learning and experimentally measured molecular properties. The absence of a conventional VL partner creates a distinct structural context for both the former VH–VL interface and the antigen-binding loops, motivating the hypothesis that HCAbs may occupy a specialized sequence space. Model-independent analyses supported this premise: functional HCAbs and conventional human VH domains showed a reproducible whole-sequence distributional distinction under source-held-out, near-duplicate- controlled analysis, and position-resolved decomposition revealed a predominantly CDR-centered organization together with a localized shift at the former VH–VL interface. The strongest compositional differences involved reduced CDR1/2 germline deviation, shorter and less hydrophobic CDR3s, and a restricted framework- interface signal rather than widespread framework remodeling. These observations provide a biological rationale for HCAb-specific pretraining while remaining agnostic as to whether individual sequence features are causal determinants of light-chain-independent folding or function.

HCAbLM converted this repertoire specialization into a distinct contextual sequence representation. Compared with conventional antibody language models, HCAbLM exhibited a region-selective compatibility prior rather than a uniform preference for HCAb sequences, with the clearest divergence occurring in CDR3. The opposing CDR3 preferences of HCAbLM and conventional antibody models persisted after controlling for CDR3-length distribution, consistent with repertoire-specific pretraining contributing to the learned sequence prior. Other regions showed more heterogeneous behavior, including framework positions whose aggregate germline- deviation frequency differed little between cohorts but whose residue contexts were less compatible with conventional antibody-model priors. Together, these results emphasize that sequence composition and contextual compatibility capture complementary aspects of antibody repertoire organization.

Frozen HCAbLM representations also retained information associated with experimentally measured developability properties. Under related-sequence-aware grouped validation, predictive performance was endpoint dependent: HCAbLM- 300M performed strongly for SEC, whereas HCAb-adapted IgBERT and ESMC-6B led the internal HIC and Tm1 benchmarks, respectively. In retrospective cross-project evaluation without refitting on project-level data, HCAbLM-300M yielded the highest observed project-macro point estimates for SEC purity and HIC favorability, whereas ESMC-6B performed best for Tm1. Because these project-level comparisons contributed to endpoint-model selection, they are interpreted as evidence of transferable and complementary representation utility rather than prospective superiority. We therefore use the term “functional grammar” in a deliberately restricted sense: sequence representations that encode information transferable to measured molecular properties, rather than evidence that the model has learned antigen specificity, affinity or biological activity in general [12,13].

More broadly, the results identify the biological composition of the pretraining repertoire as an important design variable for antibody foundation models. Specialization was most evident where the HCAb repertoire itself showed distinctive regional organization, whereas broad protein pretraining remained advantageous for thermal stability, a property governed by constraints extending beyond antibody-format-specific sequence space. These findings argue against assuming that either maximal repertoire breadth or maximal specialization will be universally optimal. Instead, the appropriate pretraining distribution is likely to depend on the molecular format and downstream property of interest. This interpretation remains associative rather than causal because HCAbLM, IgLM and AbLang-2 differ not only in training corpus but also in model architecture, training objective, scoring formulation and prior sequence exposure, and HCAbLM deliberately emphasizes CDR3 and germline- deviating residues during pretraining [14,15].

Several considerations define the scope of these conclusions. The HCAb–human VH comparisons are observational and therefore identify format-associated sequence organization rather than the causal basis of autonomous folding; Dataset B provides a provenance sensitivity analysis rather than an independent biological replication. Likewise, the four prespecified former VH–VL-interface positions represent a restricted sequence proxy for this structural surface rather than a complete contact map, and the localized signal should not be interpreted as a universal engineering rule. Finally, the retrospective developability cohorts are modest in size and selection-affected, so prospective evaluation in independently locked projects will be important for defining the generality of representation transfer. Within these boundaries, the convergent whole-sequence, position-resolved, language-model and experimental analyses support a region-selective sequence grammar for fully human HCAbs and show that repertoire-specific representation learning captures information relevant to antibody developability.

## Methods

### HCAb repertoire and unified training corpus

The starting dataset comprised HCAb repertoire-sequencing samples derived from 73 independently immunized Harbour Mice with 8 distinct antigens, generated using the fully human HCAb transgenic platform [9,10]. 6–8-week-old female mice were used and maintained under specific-pathogen-free conditions. Depending on the immunogen, mice were immunized with purified protein or antigen-expressing mammalian cells according to established protocols. Protein immunogens were assessed for identity and purity before use and were administered intraperitoneally with complete Freund’s adjuvant for the primary immunization and Ribi adjuvant for subsequent boosts; cell-based immunizations used freshly prepared antigen-expressing cells without adjuvant. Boost immunizations were performed at approximately 2–4-week intervals, with immune responses monitored by test bleeds and antigen-specific ELISA and/or flow cytometry. Animals showing suitable antibody responses received a final boost with immunogen in PBS or saline 3–5 days before lymphocyte collection. All animal procedures were performed in accordance with institutional animal-care requirements.

Single-cell B-cell receptor (scBCR-seq) libraries were generated using the sCircle Single Cell Full-Length Immunoreceptor Library Kit (Singleron) according to the manufacturer’s instructions. Briefly, isolated cells were resuspended in cold PBS at approximately 3×10^5^ cells/ml and loaded into a microwell microchip, followed by barcode beads. Cells were lysed in situ and released mRNA was captured on the barcoded beads. Retrieved beads were subjected to reverse transcription at 42 °C for 90 min, followed by cDNA amplification. A portion of the purified cDNA was retained for single-cell RNA-sequencing library preparation, whereas the remaining material was used for immunoreceptor sequencing. Full-length BCR libraries were generated through cDNA circularization, digestion and purification, followed by three rounds of BCR enrichment, fragmentation, adapter ligation, amplification, purification and size selection. Final libraries were sequenced on an Illumina NovaSeq 6000 platform.

### Data preprocessing and corpus construction

Across the repertoire datasets included in the auditable raw-read inventory, 193,405,094 paired-end heavy-chain reads entered sequence processing. Reads were quality filtered using fastp v0.23.2 and merged using FLASH v1.2.11, yielding 156,871,528 full-length contigs. Within-sample nucleotide dereplication yielded 108,584,746 unique nucleotide sequences. V(D)J assignments were generated using IgBLAST v1.22.0 [16] and restricted to productive rearrangements. Following conceptual translation, removal of C-terminal expression-tag sequence, antibody numbering and structural-validity filtering, 54,462,360 full-length VH amino-acid sequences remained. Global exact amino-acid deduplication across samples produced 36,612,172 unique candidate sequences before isolation of evaluation and holdout sets. Subsequent exclusion of evaluation-associated sequences and related sequence groups, as described below, yielded the final pretraining corpus of 31,822,392 unique HCAb VH sequences. Counts derived from annotation-support records were retained separately for provenance and label harmonization; because multiple annotation records could support the same standardized amino-acid sequence, these records were not interpreted as being directly equivalent to sequence counts in the raw-read processing cascade.

Each amino-acid position was annotated according to its relationship to the assigned V- or J-germline sequence. Reliably aligned positions matching the assigned germline residue were classified as germline matched, whereas reliably aligned positions containing a different amino acid were classified as non-germline-like (NGL). CDR3 and junctional positions, V–D–J alignment gaps, positions lacking reliable germline coverage, non-standard residues and other incompletely resolved positions were retained as a separate unresolved class. All positions were initially assigned to the unresolved class and were reclassified as germline matched or NGL only when reliable V/J alignment coverage was available outside the CDR3/junction mask. CDR3/junction annotations took precedence over J-derived framework coverage. These labels were used to define residue classes for masking and model training and should not be interpreted as direct measurements of nucleotide-level somatic hypermutation.

Exact-sequence identity was defined as identical full-length VH amino-acid sequences, whereas exact CDR3 identity was defined as identical coordinate-defined CDR3 amino-acid sequences. A near-duplicate relationship was defined between two non-identical sequences when their coordinate-defined CDR3s satisfied the prespecified sequence-similarity criterion of at least 80% amino-acid identity and at least 80% alignment coverage in both directions. For short CDR3s, for which percentage-based alignment criteria are less stable, a prespecified global edit-distance rule was applied. Near-duplicate components were defined as connected components under these pairwise relationships; thus, sequences linked directly or transitively by a qualifying near-duplicate edge were assigned to the same component. Exact full-length VH or exact-CDR3 matches were treated as related irrespective of the near-duplicate criterion.

To minimize sequence leakage between pretraining and downstream evaluation, sequence isolation was performed hierarchically at the exact-sequence, CDR3 and clone-related levels. For the designated validation and test holdouts, CDR3 similarity was assessed using a prespecified near-CDR3 criterion requiring both sequence identity and bidirectional coverage of at least 0.80; short CDR3 sequences were evaluated using a frozen global edit-distance rule. Exact full-length VH identity and exact coordinate-defined CDR3 identity were additionally used to connect related sequences into components, and complete components were assigned atomically to validation or test partitions. Pretraining candidates sharing an exact full-length VH sequence, an exact CDR3 sequence or a qualifying near-CDR3 relationship with the final holdout sequences were removed from the training pool. This step excluded 4,626,013 candidates from the 36,612,172-sequence globally deduplicated pool, leaving 31,986,159 training candidates together with validation and test sets of 80,123 and 75,634 sequences, respectively. No exact full-length VH or exact-CDR3 overlap remained between the training pool and these holdout partitions.

Additional clone-aware isolation was then applied to the frozen internal evaluation panels used for model development and assessment. Sequences were grouped using canonical clone assignments together with project/source and source-sample metadata, and complete related-sequence groups were retained within a single partition. Removal of clone-associated holdout groups and all exact amino-acid duplicates shared with the evaluation panels excluded a further 163,767 candidates, yielding the final pretraining corpus of 31,822,392 sequences. In the remaining internal evaluation panels, near-sequence similarity was used for stratification and reporting but was not used to construct additional transitive clusters or to remove further sequences from the pretraining corpus. The frozen validation partition used for model-size and checkpoint selection was defined independently of Dataset A, Dataset B and all developability outcomes; these downstream datasets did not contribute to model selection.

The final training cache stored the standardized VH amino-acid sequence, residue-level germline/NGL annotations, CDR3 coordinates, clone-group assignment, aggregated read support, V/J annotations, source metadata and a stable sequence identifier. Sequences were encoded using the native ESMC tokenizer with beginning-of-sequence and end-of-sequence tokens; residue-level CDR3 and germline-status masks were offset accordingly to maintain exact alignment between amino-acid positions and model tokens. During full-corpus pretraining, globally unique amino-acid sequences were sampled uniformly without replacement within each corpus pass. Read abundance, clone size and the number of source observations supporting a sequence were retained as metadata but were not used to replicate sequences or weight the masked-language-model objective.

### HCAb language-model training and operating-point selection

Training and evaluation partitions were defined using clone-aware hard groups. Reliable clone identifiers were scoped within project and sample namespaces; where such identifiers were unavailable, exact full-length VH amino-acid identity was used as the grouping criterion. No lineage trees were inferred. The final full pretraining corpus contained 31,822,392 unique HCAb VH sequences distributed across 6,495,575 hard groups. The frozen validation panel used for model-size and checkpoint selection contained 12,729 sequences representing 8,000 clone groups. Dataset A, Dataset B and the developability datasets were derived independently from antibody-engineering records rather than from the repertoire-sequencing corpus, and conventional human VH sequences were never included in HCAb pretraining. Evaluation HCAbs and their identified clonal relatives were excluded from the pretraining corpus according to the frozen sequence-isolation rules described above. For one designated stringent holdout, near-CDR3 relationships were defined using both sequence identity and bidirectional coverage of at least 0.80, with a prespecified edit-distance rule for short CDR3s; in the remaining internal evaluation panels, near-neighbour similarity was used for stratification and reporting rather than for additional global sequence removal.

HCAb language models were trained de novo using an ESMC-style encoder-only Transformer architecture [15], without initialization from pretrained ESMC weights. Four nominal model-size classes at 35M, 150M, 300M and 600M were evaluated. For HCAbLM-300M, the selected pass-4 checkpoint had consumed 15,200,945,920 amino-acid tokens and an estimated 3.339 × 10^19 training FLOPs. Two independent training seeds were evaluated at this operating point, and Fig. 4 reports their matched-pass diagnostic performance at the metric level rather than averaging model weights. The checkpoint carried forward for all subsequent HCAbLM representation analyses was fixed before Dataset A, Dataset B or developability outcomes were examined. Training used NVIDIA H20 GPUs, with increasing accelerator allocation for larger model configurations; estimated compute in Fig. 4 was calculated as *C* « *6ND* , where *N* is the non-embedding parameter count and *D* is the number of consumed amino-acid tokens.

Residue-level training labels were derived from V- and J-germline alignments in the pretraining corpus and were distinct from the allele-resolved IGHV annotation used for the HCAb–human VH repertoire comparisons. Reliably aligned positions were classified as germline matched when the observed amino acid agreed with the assigned germline residue and as non-germline-like (NGL) when it differed. CDR3 and junctional positions, V–D–J alignment gaps, uncovered positions, ambiguous germline assignments and other incompletely resolved sites were retained as a separate unresolved class. True CDR3 positions were defined independently from sequence coordinates and were not assigned a germline/NGL state.

Masked-language-model pretraining used a fixed masking budget of 15% of amino-acid positions per sequence. Positions belonging to either the true CDR3 or the high-confidence NGL class outside CDR3 were preferentially sampled with probability 0.25. If the sampled priority positions exceeded the total masking budget, they were randomly subsampled to the budget; otherwise, the remaining masked positions were drawn uniformly from background residues. Selected positions were corrupted according to an 80:10:10 scheme, with 80% replaced by the mask token, 10% replaced by a random amino acid and 10% left unchanged.

Model optimization used a region-balanced focal-loss objective. For masked tokens belonging to residue class *r*,

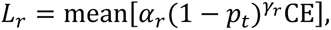

where *p_t_* is the model probability assigned to the observed amino acid and CE is the token-level cross-entropy.

Full-sequence validation used position-wise single-mask pseudo-cross-entropy (PCE), averaged at the clone level to reduce disproportionate contribution from expanded sequence families. Let PCE_NGL_denote the mean PCE at high-confidence NGL positions outside CDR3 and PCE_CDR3_the corresponding value at true-CDR3 positions. The prespecified regional validation summary was

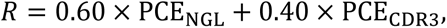

with regional predictive score

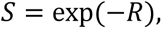

such that higher values indicate lower region-weighted prediction loss. Training trajectories were monitored using a frozen diagnostic bank comprising 80,000 masked target positions sampled equally from four strata: 20,000 germline-matched non-CDR3 positions, 20,000 high-confidence NGL non-CDR3 positions, 20,000 unresolved or other non-CDR3 positions and 20,000 true-CDR3 positions. Stratum-level estimates were combined using inverse-probability weighting to recover the positional mixture of the underlying validation data. Diagnostic-bank PCE was used to visualize training trajectories and compare performance–compute behavior, whereas final model-size and checkpoint selection was based on the frozen full-sequence validation panel. Neither the independent test panels, Dataset A, Dataset B nor any developability outcome contributed to model-size or checkpoint selection. HCAbLM-300M reached the observed validation-score plateau with less estimated compute than the 600M configuration and was therefore selected as the performance–compute operating point used in subsequent analyses.

### HCAb adaptation of comparator models

HCAb-adapted IgBERT and ESMC-LoRA were generated by low-rank adaptation (LoRA) [20] of pretrained encoders on HCAb corpus. Adaptation combined 60% clone-balanced sampling with 40% row-uniform sampling. The masking scheme mixed standard masking (45%) with region-targeted masking (55%). Both adaptations used AdamW, a 500-step learning-rate warm-up followed by cosine decay, weight decay 0.01 and global gradient-norm clipping at 1.0.

HCAb-adapted IgBERT was initialized from Exscientia/IgBert. LoRA adapters targeted the query, key, value and attention-output projections in the final 10 Transformer layers, with rank 16, scaling parameter α = 32 and dropout 0.05, giving approximately 1.31 million trainable parameters. The peak learning rate was 1 × 10⁻⁵, and the objective was sequence-normalized masked-token cross-entropy. The training run continued for five epochs (27,700 steps; approximately 5.6 h on four NVIDIA H20 GPUs).

HCAb-adapted ESMC-LoRA was initialized from EvolutionaryScale/esmc-600m-2024-12. LoRA adapters targeted the QKV and attention-output projections in the final 24 Transformer layers, with rank 8, scaling parameter α = 16 and dropout 0.05, giving approximately 1.33 million trainable parameters. The peak learning rate was 1 × 10⁻⁵. Training lasted two epochs (11,080 steps; approximately 1.64 h on eight NVIDIA H20 GPUs) and used masked-token loss plus a germline-status auxiliary loss weighted by 0.05.

Downstream partitions kept exact-sequence, clone/parent and near-duplicate groups intact. However, the adaptation corpus was not globally purged of sequences associated with all later benchmarks before adaptation. Whole-exposure-group exclusion was therefore additionally evaluated as a sensitivity analysis. These adapted comparators should not be interpreted as having the same pretraining-exposure exclusions as HCAbLM.

### HCAb–human VH cohorts and repertoire analysis

Dataset A comprised 4,839 exact amino-acid-sequence-unique VH domains containing only standard amino acids: 1,429 mature functional fully human HCAbs and 3,410 publicly available assay-positive human VH sequences. The HCAb cohort represented 17 independent source units and consisted of molecules that had undergone functional validation within antibody-discovery campaigns. Because functional evidence in the internal and public datasets was generated using heterogeneous assays and was not harmonized to a common potency threshold, the two cohorts are described as mature functional HCAbs and public assay-positive human VH sequences without implying equivalence of the underlying functional assays.

The public human VH cohort was assembled from CoV-AbDab, IEDB and CATNAP [22–24] and frozen for analysis on 7 August 2026. The CoV-AbDab component was based on the 8 February 2024 snapshot, the CATNAP component on the 1 August 2026 release, and IEDB records were retrieved through the application programming interface available at the time of dataset construction. After resource-specific eligibility filtering, 3,693 candidate sequences were reviewed, of which 283 were excluded and 3,410 retained. Because some antibody sequences were represented in more than one resource, exact amino-acid deduplication was performed across databases before analysis. The retained public cohort represented 360 source units, defined at the level of the smallest independently identifiable study or sequence collection; sequences derived from the same donor or directly related collection were assigned to the same source unit.

Eligibility criteria were adapted to the provenance information available in each public resource. IEDB records were required to encode a complete human heavy-chain protein sequence, originate from a human antibody lineage and have a positive qualitative assay annotation. CATNAP records were required to derive from human donors and to have associated neutralization evidence, whereas CoV-AbDab records were required to have documented human origin together with binding or neutralization evidence. Direct human B-cell lineages, human-donor repertoire-derived antibodies and optimized descendants of a documented human parent were retained. Animal-derived humanized or CDR-grafted antibodies, de novo or unresolved display-library sequences, and records lacking sufficient evidence of human lineage were excluded. Record-level provenance, including database identifiers, source publications and available assay metadata, was retained to support source assignment and auditing.

Because the HCAb transgenic platform encodes a restricted human IGHV repertoire, conventional human comparators were restricted to the same supported IGHV gene set rather than sampled from unrestricted human VH sequence space. The supported set comprised IGHV3-11, IGHV3-23, IGHV3-30, IGHV3-33, IGHV3-48, IGHV3-53, IGHV3-64, IGHV3-66, IGHV3-74 and IGHV6-1. Inclusion required an unambiguous gene-level assignment to one of these genes by IgBLAST. This restriction aligned the set of germline genes represented in the two cohorts but did not equalize their usage frequencies; analyses were therefore repeated with equal weighting across shared IGHV strata as a sensitivity analysis. No one-to-one sequence matching between HCAb and human VH molecules was performed. The position-resolved Dataset A analyses used the full supported IGHV set, whereas the primary whole-VH distributional analysis was further restricted to the five IGHV genes with sufficient independent source representation in both cohorts.

Dataset B comprised 232 exact-sequence-unique HCAbs and 232 exact-sequence-unique conventional H2L2 human VH domains from a separate set of functionally characterized antibodies and was analyzed independently as a provenance sensitivity cohort. The cohorts were exactly balanced across the three represented IGHV genes: IGHV3-33, 111 HCAbs and 111 human VH sequences; IGHV3-23, 79 and 79; and IGHV3-30, 42 and 42. Related sequences were grouped into frozen near-duplicate components, and each component contributed equal total weight to the analysis. Dataset B was not pooled with Dataset A. Because independent source-level provenance and molecular ancestry could not be reconstructed to the same resolution as in Dataset A, Dataset B was used as a component-level sensitivity analysis rather than as an independent source-level replication.

### Cross-fitted whole-VH distribution analysis

The primary whole-VH analysis was performed on Dataset A after removal of 35 sequences belonging to cross-cohort near-duplicate components. To ensure adequate source-level replication, analysis was restricted to IGHV genes represented by at least five independent source units in each cohort. Five genes met this criterion—IGHV3-11, IGHV3-23, IGHV3-33, IGHV3-53 and IGHV3-74—yielding 2,945 sequences comprising 1,400 HCAbs from 17 source units and 1,545 human VH sequences from 184 source units. IGHV3-30, IGHV3-48 and IGHV3-66 were excluded from this primary analysis because the HCAb cohort contained only one, four and three source units, respectively.

Each complete VH sequence was represented using two complementary, model-independent feature sets. First, IMGT-numbered sequences were encoded by residue or gap identity across the 155 IMGT coordinates observed in the eligible pooled dataset, including insertion positions. Second, complete-sequence composition was represented by amino-acid 2-mer and 3-mer spectra, comprising 8,400 features. Together, these representations yielded 11,655 sequence features. Cosine similarity was calculated independently for the aligned-position and k-mer representations, and the prespecified whole-VH kernel was defined as their equal-weight average. No HCAbLM embedding or other representation pretrained on the HCAb repertoire was used in this analysis.

Fivefold cross-fitting was performed separately within each IGHV stratum at the source-unit level. All sequences from a given source were assigned to the same fold, and each sequence received a score only when its source was held out. Within each fold, training sequences belonging to a frozen near-duplicate component represented in the held-out fold were removed, leaving no shared near-duplicate components between training and test data. For a held-out sequence *X*, the witness score was defined as

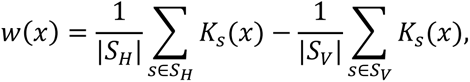

where *k* denotes the prespecified whole-VH kernel. For each retained training source s, K_s_(X) is the mean kernel similarity between x and the sequences from that source. *S_H_* and *S_v_* denote the HCAb and human VH training-source sets remaining after fold-specific exclusion. Positive values therefore indicate greater similarity to the HCAb training distribution and negative values greater similarity to the human VH distribution. The primary cross-fitted witness separation was calculated as the difference between mean out-of-fold witness scores for HCAbs and human VH sequences, with equal weight assigned to each of the five IGHV strata. This statistic is a cross-fitted two-sample separation measure in raw kernel units and is distinct from a conventional in-sample *MMD^2^* .

For visualization and source-level comparison, witness scores were standardized within each IGHV-by-fold block using the distribution of training-source means. Density estimates assigned equal total weight to each IGHV stratum, each source within an IGHV stratum and each sequence within a source. Uncertainty in the primary separation statistic was estimated using 5,000 source-level bootstrap replicates applied to the fixed out-of-fold scores. Statistical significance was assessed using 5,000 source-level label permutations within IGHV-by-fold blocks. Four of the five IGHV-specific tests reached the minimum attainable permutation *P* value of 1/5,001 , and IGHV3-74 yielded *P* = 0.0028 ; all five remained significant after Holm correction. The global-null significance reported in Fig. 2 was conservatively summarized using a Bonferroni bound across the five prespecified IGHV-specific tests. Source-level visualization in Fig. 2b used one mean standardized out-of-fold witness score per source unit.

As a complementary discrimination analysis, an L2-regularized logistic classifier was fitted separately within each IGHV stratum using the same whole-sequence features, source-held-out folds, fold-specific near-duplicate exclusion and weighting hierarchy. Classifier scores were averaged within source before calculating source-level AUROC, and the primary summary was the equally weighted mean across the five IGHV-specific AUROCs. This yielded a mean source-level AUROC of 0.976, with a 95% source-bootstrap confidence interval of 0.955–0.992. Out-of-fold sigmoid outputs were treated as uncalibrated discrimination scores rather than probabilities of HCAb class membership. The cross-fitted kernel witness analysis was prespecified as the primary model-independent two-sample test, with the classifier serving as complementary evidence for separability of the two whole-VH sequence distributions.

### Germline annotation, regional composition and statistical analysis

For cohort comparisons, and separately from the V/J labels used during pretraining, sequences were annotated with ANARCII v2.0.8 using IMGT coordinates [18,19]. Within reliably mapped V-derived positions in FR1, CDR1, FR2, CDR2 and FR3, an observed amino acid differing from the mapped IGHV germline residue was classified as NGL; identical residues were classified as germline matched. CDR3, including insertion positions, was analyzed as a separate class and was not assigned an NGL state. FR4 was excluded from the primary NGL partition because allele-resolved IGHJ assignments were unavailable. Positions for which ambiguous allele assignments implied different germline residues, V-region insertions and unmappable coordinates were classified as unknown and excluded from NGL numerators and denominators. Training-corpus labels used the frozen numbering and region-boundary fields carried in the sample-level reports, whereas cohort analyses were independently re-annotated with ANARCII v2.0.8; the two coordinate streams were not mixed within an analysis.

Model-independent repertoire analyses included occupied length in each of the seven IMGT regions; regional fractions of each of the 20 amino acids; and six prespecified physicochemical classes: acidic (D,E), basic (H,K,R), aromatic (F,W,Y), polar uncharged (C,N,Q,S,T), aliphatic/hydrophobic (A,I,L,M,V) and turn-special (G,P). For V-derived regions, we additionally calculated NGL fraction, observed NGL-residue burden and germline-to-observed substitution burden. Because amino-acid and residue-class fractions are compositional, individual components were interpreted as marginal cohort differences rather than independent mechanistic effects.

### Former VH-VL interface analysis

The V-derived framework was defined as standard amino-acid residues at integer IMGT positions within FR1 (1–26), FR2 (39–55) and FR3 (66–104). A restricted former VH–VL-interface hallmark patch was prespecified as IMGT positions 42, 49, 50 and 52, with all remaining callable framework positions forming the comparison partition. This four-position set was used as a sequence-level proxy for the former VH–VL interface and was not intended to represent a complete structural contact map.

Primary estimands were the source-equal HCAb–human VH difference in NGL fraction within the hallmark patch and remaining framework, together with the patch-by-cohort interaction, defined as the difference between these two cohort effects. Sequence-level measurements were first aggregated within source, and source units contributed equal total weight within each cohort. Uncertainty was estimated using 5,000 two-stage bootstrap replicates in which source units were resampled first and frozen near-duplicate components were resampled within source. Pointwise 95% confidence intervals were obtained from the centered bootstrap distribution. Benjamini–Hochberg false-discovery-rate adjustment was applied within prespecified comparison families [28], with an adjusted q ≤ 0.05 and a pointwise confidence interval excluding zero required for a primary difference call. Analyses additionally required at least 80% source coverage in each cohort.

To determine whether the interface-associated signal was robust to repertoire composition, the primary analysis was repeated with equal weighting across shared IGHV strata and in leave-one-HCAb-source-out analyses. Logit-scale analyses provided an additional sensitivity assessment. For physicochemical composition, residue-class fractions were calculated separately for the hallmark patch and remaining framework within each cohort, and the HCAb–human VH difference in the resulting patch-versus-framework contrast was evaluated using the same source-aware resampling framework.

An exploratory position-resolved analysis was performed across the 96 integer IMGT V-derived positions callable in both cohorts. Position-level effects were summarized as source-equal HCAb–human VH differences in NGL frequency. A position was considered to show a pointwise signal only when its 95% confidence interval excluded zero and its within-family Benjamini–Hochberg-adjusted qvalue was ≤ 0.05. To distinguish localized signals from stronger position-specific evidence, family-wise simultaneous confidence intervals were additionally calculated using the maximum absolute centered bootstrap deviation across positions. Positions meeting the pointwise criterion but not the simultaneous-interval criterion were interpreted as localization signals rather than confirmatory single-residue effects. These analyses therefore test for a localized repertoire association at the prespecified former VH–VL-interface patch and do not establish a causal adaptation mechanism or a sequence-design rule.

### Language-model scoring and cross-model comparison

Sequence-context compatibility in Dataset A was evaluated using frozen HCAbLM-300M, IgLM [5] and AbLang-2 [14] models. For HCAbLM-300M, each residue was masked individually within the complete VH sequence and the model probability assigned to the observed amino acid was recorded after normalization over the 20 standard amino acids. Residue-level pseudo-cross-entropy (PCE) was defined as the negative logarithm of this probability. IgLM was evaluated using its frozen checkpoint with heavy-chain and human-species conditioning and its native bidirectional infilling formulation [5]. AbLang-2 was evaluated using the frozen public checkpoint and bidirectional masked-token scoring [14]. All models were evaluated without task-specific fine-tuning on Dataset A.

Because HCAbLM, IgLM and AbLang-2 differ in architecture, training objective and scoring formulation, their raw prediction losses were not assumed to share a common numerical scale. Cross-model comparisons were therefore based on standardized cohort effects rather than direct comparison of raw losses. For each model and prespecified sequence region, the HCAb–human VH difference in mean prediction loss was divided by the pooled standard deviation of source-unit mean prediction loss. This standardization uses between-source variation as the reference scale and therefore does not treat individual residues as independent observations. Cohort effects were defined as HCAb minus human VH; negative values indicate lower prediction loss, and thus greater sequence compatibility, for HCAbs under the corresponding model.

To assess whether CDR3 prediction-loss differences could be explained by differences in CDR3-length distributions between cohorts, source units were assigned equal weight within each integer CDR3-length stratum represented in both cohorts, after which the 18 shared length strata were weighted equally. This length-balanced analysis was treated as a descriptive sensitivity analysis.

Uncertainty in standardized effects was estimated using the same source-aware resampling structure applied to the Dataset A repertoire analyses. Cross-model contrasts were calculated within shared bootstrap replicates because all models were evaluated on the same sequences and source units. CDR3 was analyzed independently of germline-deviation status because germline/NGL annotation was defined only for reliably mapped V-derived positions. Alternative autoregressive scoring of IgLM was evaluated as a sensitivity analysis and is reported in the figure Source Data.

### Developability datasets and experimental measurements

Developability data were obtained from internal HCAb antibody-engineering projects that were not part of the repertoire-sequencing corpus. Missing observations were excluded from the corresponding analysis and no imputation was done on any missing values.

Size-exclusion chromatography (SEC) was performed using an Agilent 1260 Infinity II HPLC system equipped with a TSKgel G3000SWxl analytical SEC column. Purified antibody samples were analyzed at a concentration of 1 mg/ml, with an injection volume of 20 µl. The mobile phase was 1× PBS, pH 7.4 (Sangon Biotech, E607016), delivered isocratically at 1.0 ml/min at room temperature. Protein elution was monitored by absorbance at 280 nm. Chromatograms were integrated using ChemStation software, and SEC purity was calculated from the integrated chromatographic peak areas as the fraction attributable to the monomeric species relative to the total integrated protein-associated peak area. For the internal prediction benchmark, SEC was formulated as a binary endpoint using a prespecified purity threshold of <85%.

Hydrophobic interaction chromatography (HIC) was performed using an Agilent 1260 Infinity II HPLC system equipped with a TSKgel Butyl-NPR analytical column (4.6 × 35 mm). Samples were analyzed at 1 mg/ml with an injection volume of 20 µl and monitored at 280 nm. Chromatographic separation was performed at a flow rate of 0.7 ml/min using a 16-min linear gradient from 100% mobile phase A (20 mM phosphate buffer, 1.8 M ammonium sulfate, pH 6.0) to 100% mobile phase B (20 mM phosphate buffer, pH 6.0). Chromatograms were integrated using ChemStation software. HIC behavior was summarized using the experimentally derived HIC index, with higher values corresponding to lower HIC favorability.

Thermal stability was measured by differential scanning fluorimetry (DSF) using SYPRO Orange dye and a Bio-Rad CFX96 real-time PCR system. Antibody samples and SYPRO Orange were combined in 96-well PCR plates (Thermo Fisher Scientific, AB-0700/W) in the assay buffer to a final volume of 40 µl per well. Plates were sealed and incubated at 25 °C for 5 min before the temperature was increased from 25 °C to 95 °C at 0.2 °C every 0.2 min (equivalent to approximately 1 °C per min). Fluorescence was monitored throughout the thermal ramp, and melting transitions were analyzed using Bio-Rad CFX Maestro software. Tm1 was defined as the temperature corresponding to the first resolved thermal-unfolding transition of the antibody sample.

### Developability prediction benchmarks and model evaluation

All sequences included in the developability benchmarks, together with identified clonal relatives, were excluded from HCAbLM pretraining before model fitting. Related-sequence groups were defined using exact full-length VH identity, recorded parent–clone relationships and prespecified near-duplicate relationships, and each connected group was assigned wholly to a single cross-validation partition. Candidate encoders were evaluated using frozen sequence representations under a common split specification. The evaluated representations comprised HCAbLM-300M, ESMC-300M, HCAb-adapted IgBERT, IgBERT, ESMC-6B, HCAb-adapted ESMC-LoRA and ESMC-600M. HCAbLM embeddings were generated using the frozen HCAbLM-300M checkpoint selected by the operating-point procedure described above; the encoder was not further trained or adapted using developability outcomes.

Endpoint-specific prediction heads were intentionally shallow to evaluate information already present in the frozen representations. SEC was modeled using an L2-regularized logistic-regression head, whereas HIC and Tm1 were modeled using L2-regularized ridge-regression heads. Hyperparameters were selected within the nested training procedure, yielding *C* = 0.01 for SEC and ridge penalties of *α =* 100 and *α* = 1,000 for HIC and Tm1, respectively. No project identity, assay-condition metadata or other experimental covariates were supplied to the prediction heads.

For internal evaluation, SEC was formulated as a binary classification endpoint identifying antibodies with purity below 85% and was evaluated using area under the receiver-operating-characteristic curve (AUROC). HIC was analyzed as a continuous endpoint after sign inversion of the measured HIC index such that higher values represented more favorable chromatographic behavior. Tm1 was analyzed as the continuous first thermal-transition midpoint. For cross-project evaluation, SEC purity was treated as a continuous measurement and ranked against the model-derived SEC score using Spearman correlation; because the internal classifier was trained to predict low SEC purity, its output was sign-inverted so that higher model scores consistently denoted more favorable SEC behavior. HIC and Tm1 were likewise evaluated using Spearman correlation.

Internal performance was estimated using nested grouped cross-validation with five outer and four inner folds. Related-sequence groups, rather than individual sequences, were used as the splitting unit, and each group contributed equal total weight to model fitting and evaluation. Feature standardization and hyperparameter selection were performed exclusively within the corresponding outer-training partition. The SEC dataset contained 1,469 sequences in 773 related-sequence groups, the HIC dataset contained 506 sequences in 246 groups and the Tm1 dataset contained 425 sequences in 209 groups.

Cross-project transfer was evaluated retrospectively using antibody projects not included in the corresponding model-fitting partitions. After strict whole-group exclusion of related sequences, the evaluation set contained 169 sequences in 94 groups. Endpoint-specific support comprised 87 sequences from two projects for SEC and 108 sequences from three projects each for HIC and Tm1. Prediction heads were not refitted using these project-level evaluation data. Spearman correlations were calculated separately within each project and then macro-averaged with equal weight across projects. Confidence intervals and paired model comparisons were estimated using 2,000 shared bootstrap resamples performed independently within each fixed project.

Seven frozen encoder representations were compared in the retrospective cross-project analysis. Because performance on these project-level data contributed to endpoint-level model comparison, the resulting estimates were considered selection-affected and were interpreted as retrospective benchmarking rather than as independent prospective validation. Accordingly, project-level results are reported as comparative point estimates with uncertainty rather than as evidence of prospective model superiority.

### Statistical analysis and reproducibility

NGS corpus characteristics, repertoire summaries and model-scaling trajectories were treated as descriptive analyses. Inferential analyses accounted for the relevant dependence structure of each dataset rather than treating individual residues or related antibody sequences as independent observations. Depending on the analysis, the statistical unit was the independent source, frozen near-duplicate component, related-sequence group or antibody project, as specified in the corresponding Methods sections. Unless otherwise stated, statistical tests were two-sided.

For Dataset A repertoire analyses, uncertainty was estimated by resampling independent source units while preserving the hierarchical structure of sequences within sources and IGHV strata. Whole-VH distributional significance was assessed by source-level label permutation within IGHV and cross-validation strata, with Holm adjustment across the five prespecified IGHV-specific tests; the global-null result was conservatively summarized using a Bonferroni bound. Position-resolved and regional analyses used source-aware bootstrap confidence intervals and Benjamini– Hochberg correction within prespecified families of related hypotheses. Where simultaneous family-wise confidence intervals were used to distinguish exploratory localization from confirmatory position-level effects, these are reported with the corresponding analyses.

Developability benchmarks used related-sequence groups as the splitting and weighting unit during internal cross-validation. For retrospective cross-project evaluation, correlations were calculated within individual antibody projects and macro-averaged with equal project weight. Confidence intervals and paired model comparisons were obtained by shared bootstrap resampling within fixed projects, thereby preserving project membership and enabling paired comparison of models on the same resampled observations.

Model-training seeds represented independent optimization runs and were distinct from seeds used for data partitioning or statistical resampling. Randomized analyses were performed using fixed seeds specified before execution to ensure computational reproducibility; exact implementation seeds are retained with the analysis records but do not affect the definition of the statistical estimands. Missing endpoint measurements were excluded from the corresponding endpoint analysis and were not imputed.

Exact sample sizes, effect estimates, confidence intervals, adjusted q values and paired model-comparison statistics are reported in the figure Source Data. No statistical method assumed independence of individual residues when inference was made at the sequence, source, related-group or project level.

## Use of generative-AI-assisted writing tools

Generative AI tools were used to assist manuscript organization and language editing. They were not used to generate experimental observations, derive the reported numerical results or execute model training. All analyses, interpretations, citations and final wording were reviewed and verified by the authors, who take full responsibility for the manuscript.

## Data availability

The fully human HCAb NGS repertoire and associated sequence-level metadata used for pretraining are proprietary to Nona Biosciences (Shanghai) Co., Ltd. and cannot be made publicly available because of intellectual-property, contractual and project-specific data-use restrictions. Dataset A also contains proprietary HCAb sequences and therefore cannot be redistributed in full.

Source data underlying Figs. 1–6 and Extended Data Figs. 1–5, including analysis-ready derived data, cohort and grouping identifiers, aggregate model outputs and the public-comparator record list, will be deposited in a persistent archival repository before publication. Public human VH comparator records were obtained from IEDB, CATNAP and CoV-AbDab [22–24] and can be accessed through the originating resources. The deposited comparator manifest will provide source identifiers, publications and dataset version or retrieval information sufficient to reconstruct the public component of the analysis.

## Code availability

Analysis and evaluation code sufficient to reproduce the reported statistical analyses and figures from the shareable frozen inputs will be deposited with environment specifications, random seeds, configuration files and artifact hashes. HCAbLM training code and model weights are proprietary to Nona Biosciences (Shanghai) Co., Ltd. and are not planned for unrestricted public release. Because the custom model and code are central to the manuscript, the authors will provide editors and referees with confidential access to the relevant checkpoint and code, or to an executable evaluation environment that permits verification of the reported model outputs, subject to intellectual-property and security restrictions.

## Ethics and permissions

Animal procedures were conducted in accordance with applicable institutional requirements. Public database records were used subject to the terms of the originating resources.

## Extended Data figures

**Extended Data Fig. 1.**
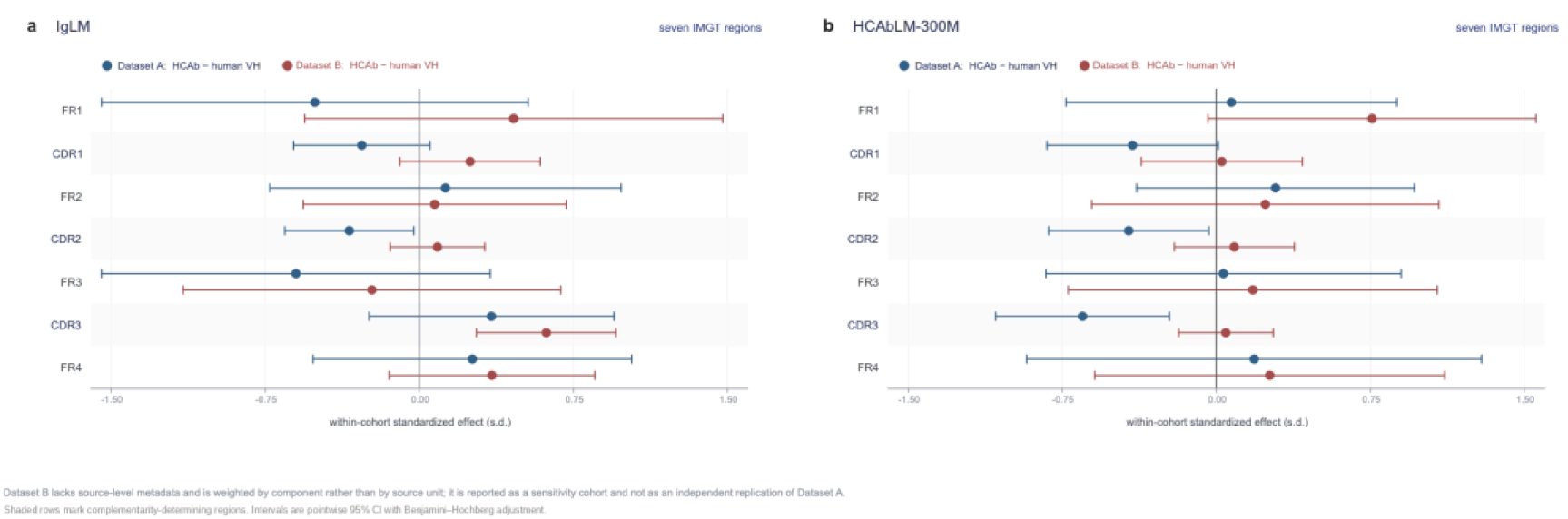
Regional prediction-loss effects in two HCAb–human VH cohorts. Within-cohort standardized HCAb - human effects are shown for **a,** IgLM and **b,** HCAbLM-300M; the corresponding AbLang-2 seven-region decomposition is provided in the Source Data. Dataset A contains 1,429 HCAbs and 3,410 human VH sequences and uses source-equal weighting; Dataset B contains 232 HCAbs and 232 in-house human VH sequences and uses near-duplicate-component-equal weighting. Whiskers indicate pointwise 95% bootstrap CIs and q values were adjusted within each seven-region cohort-by-model family. The reproducible feature across cohorts is regional non-uniformity, not the sign of every individual regional or whole-domain effect.

**Extended Data Fig. 2.**
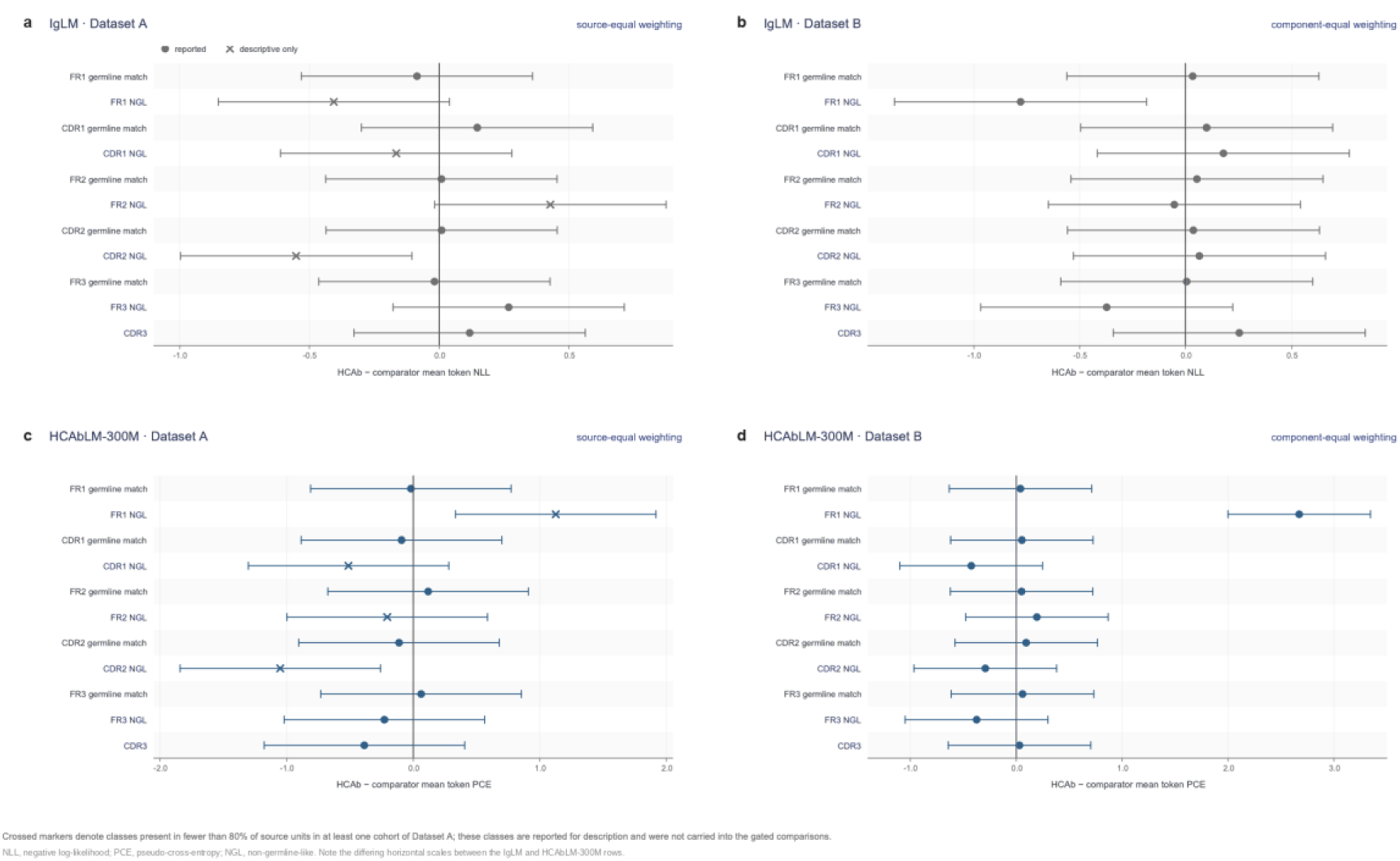
Region-by-germline-status decomposition of prediction-loss effects. HCAb - human differences in per-residue prediction loss are decomposed by IMGT region and germline-match/NGL status, with CDR3 analyzed separately, for IgLM and HCAbLM-300M in Datasets A and B. Dataset A uses source-equal weighting and Dataset B near-duplicate-component-equal weighting. Whiskers indicate pointwise 95% bootstrap CIs; q values were adjusted across the prespecified eleven-class family. Crosses identify classes represented in <80% of source units in at least one Dataset A cohort and are interpreted descriptively. IgLM and HCAbLM scores are shown on their native axes.

**Extended Data Fig. 3.**
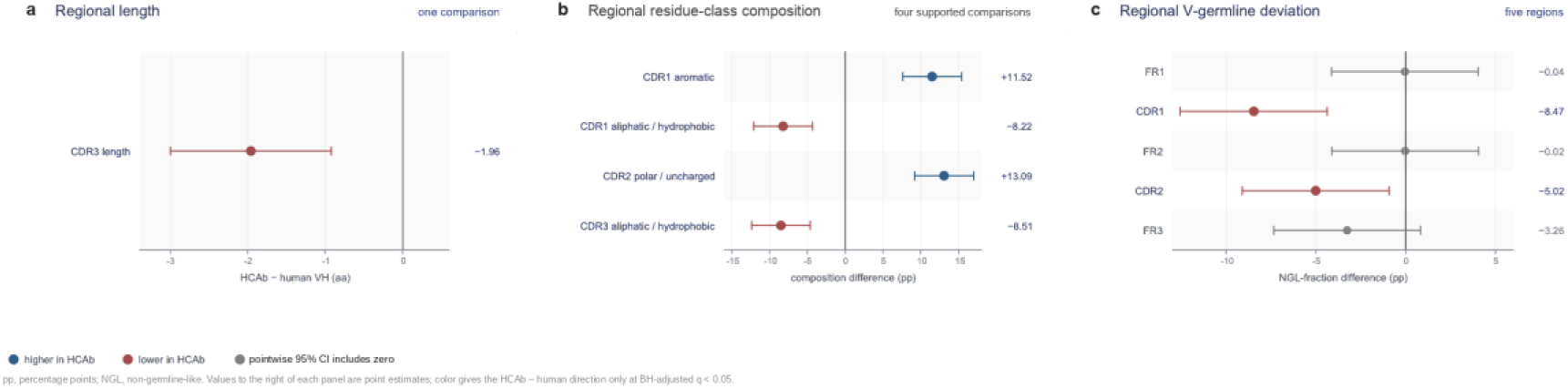
Model-independent HCAb–human VH repertoire differences are concentrated in CDRs. Source-equal HCAb - human effects in Dataset A are shown for **a,** CDR3 length; **b,** physicochemical residue-class fractions meeting q < 0.05; and **c,** regional NGL fractions at reliably mapped V-derived positions. Points indicate source-equal estimates and whiskers pointwise 95% bootstrap CIs. Color indicates effect direction only for comparisons meeting the prespecified q < 0.05 and source-coverage criteria. Leave-one-HCAb-source-out and shared-IGHV-equal-weighting sensitivity analyses are reported in Source Data.

**Extended Data Fig. 4.**
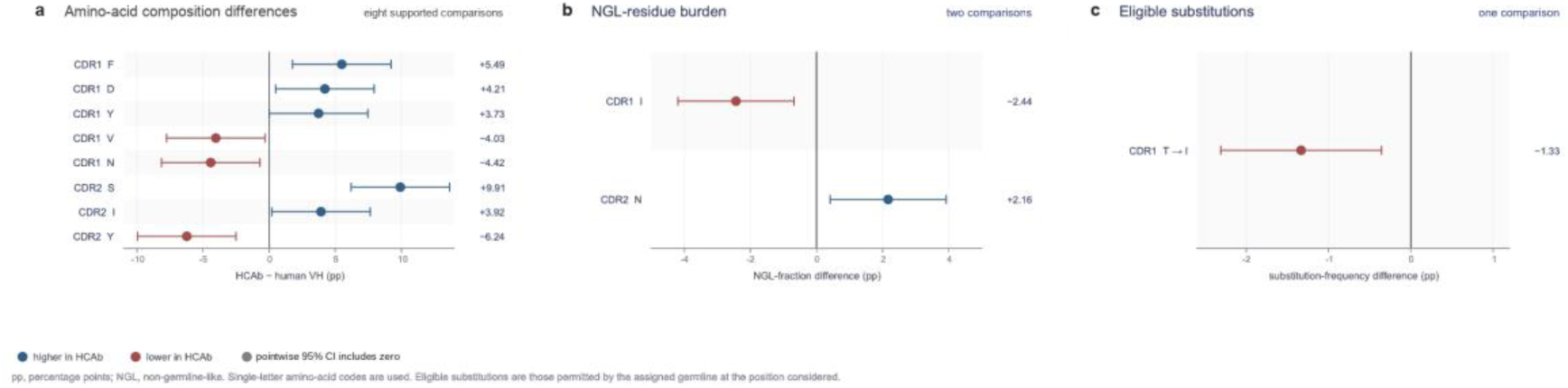
Supported amino-acid-level repertoire differences are confined to CDR1/2. **a,** Regional amino-acid fractions meeting the prespecified q < 0.05 criterion; each comparison concerns the fraction of one amino acid within a region and is not a single-IMGT-position test. **b,** Supported differences in NGL observed-residue burden. c, The eligible germline-to-observed substitution meeting the same criterion. Substitution eligibility was frozen before group comparison and required ≥20% source coverage and ≥20 observations in each cohort. Points show source-equal HCAb - human estimates and whiskers pointwise 95% bootstrap CIs.

**Extended Data Fig. 5.**
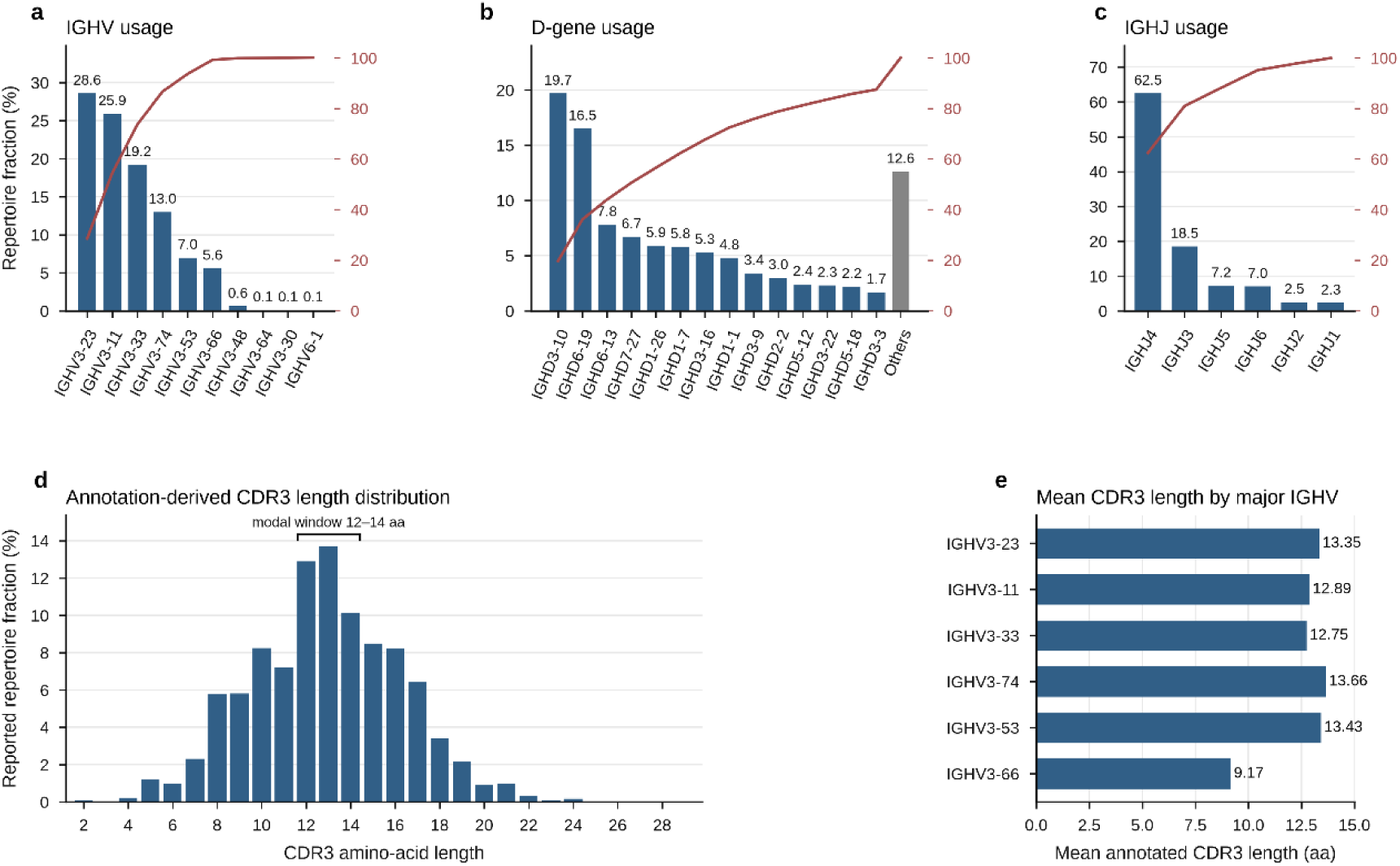
V(D)J gene use and annotation-derived CDR3 length in the HCAb pretraining corpus. Panels summarize the 31,822,392 globally exact-amino-acid-unique full training sequences, assigned to 6,495,575 hard groups, and the callable fraction for each annotation. **a–c,** IGHV, IGHD and IGHJ gene distributions, with cumulative fractions after genes were ordered by decreasing abundance; low-frequency IGHD genes are pooled as ‘Others’. **d,** Annotation-derived CDR3 amino-acid length distribution; the bracket marks the modal 12–14-amino-acid window. **e,** Mean annotation-derived CDR3 length for the six most abundant IGHV genes. These corpus-level distributions are distinct from Dataset A/B and should not be interpreted as the IGHV composition of the comparison cohorts.

